# A genetic signature predicts aggressive paraganglioma sensitivity to dual PI3K-CDK4/6 inhibition therapy

**DOI:** 10.1101/2025.02.18.638668

**Authors:** Bhargavi Karna, Federica Gentile, Sebastian Gulde, Daniela De Martino, Julian Maurer, Katyayni Ganesan, Katharina Wang, Nicole Bechmann, Elena Rapizzi, Mercedes Robledo, Ester Arroba, Stephan Herzig, Svenja Nölting, Hermine Mohr, Natalia S. Pellegata

## Abstract

Effective medical therapies for metastatic paraganglioma (mPPGL) are currently lacking, leading to dismal prognosis. Building on our knowledge of molecular mechanisms driving PPGL progression, we assessed the therapeutic potential of targeting two critical processes: PI3K signaling and cell cycle regulation. The efficacy of buparlisib (PI3Ki) and ribociclib (CDK4/6i), individually and combined, was assessed *in vitro* using PPGL cell lines, rat– and patient-derived primary cells, and *in vivo* using PPGL cells-derived mouse xenografts. The combination therapy demonstrated superior antitumor activity compared to single agents, particularly *in vivo*. Mechanistically, the efficacy of the combination therapy was associated to the downregulation of FOXM1-controlled genes implicated in mitotic spindle assembly and chromosomal segregation, leading to mitotic catastrophe. Data mining and qRT-PCR showed this genetic signature to be upregulated in human mPPGLs. This suggests that aggressive PPGLs exhibit heightened vulnerability to dual PI3K and CDK4/6 inhibition, offering a promising therapeutic avenue for these challenging cancers.

## Introduction

Paragangliomas (PPGL) are rare neuroendocrine tumors arising from chromaffin cells in the adrenal medulla (pheochromocytomas) or parasympathetic paraganglia [1]. While typically benign, 10–15% of pheochromocytomas and 35–40% of extra-adrenal paragangliomas become metastatic, with 11–31% of cases presenting with metastases at diagnosis [2,3]. Surgery is the only curative treatment for localized PPGL, but it is often unfeasible in metastatic cases [4]. For slow-progressing metastatic (m)PPGL, radionuclide therapy ([177Lu]-DOTA-TATE or [131I]-MIBG) achieves disease stability in about 50% of patients [5,6], while rapidly progressing cases are treated with chemotherapy (e.g., CVD or temozolomide ± capecitabine), though response rates are below 40%, with no proven survival benefit [7–9]. Emerging targeted therapies, including sunitinib, showed limited efficacy [10]. In addition to the lack of effective treatment, the clinical management of patients with aggressive PPGLs is further complicated by the absence of reliable markers predictive of metastatic risk. These limitations lead to the poor prognosis associated with mPPGL.

PPGLs are mainly sporadic but can be hereditary [11,12]. Next generation sequencing (NGS) revealed that over 30-40% of PPGL patients carry germline mutations in at least 21 cancer susceptibility genes, with an additional 40-50% of patients bearing known somatic driver-mutations [11,12]. Based on transcriptome, catecholamine secretion and clinical behavior, PPGLs are sub-grouped in different molecular subtypes: a pseudohypoxic, a kinase signaling, and a Wnt-activated cluster [11,12]. The pseudohypoxic cluster (Cluster 1) includes tumors with mutations in genes regulating hypoxia signaling pathways (e.g. *SDHx*, *FH*, *VHL*, *EPAS1*, among others). Pseudohypoxic PPGLs are the most aggressive tumors [13,14], and over 90% of the pediatric cases belong to this cluster [15]. Cluster 2 comprises PPGLs with mutations in *RET, NF1, KIF1B*, *TMEM127*, and *MAX* genes, which are associated with abnormal activation of phosphatidylinositol 3′ kinase (PI3K)/protein kinase B (AKT), mitogen-activated protein kinase (MAPK)/extracellular signal-regulated kinase (ERK), and mammalian target of rapamycin C1 (mTORC1)/p70S6K [12,16]. PPGLs with *CSDE1* mutations or *MAML3* gene fusions belong to the Wnt-altered Cluster 3 [11,12].

Overactivation of the PI3K/AKT/mTOR pathway has been found in hereditary as well as sporadic PPGLs, caused by e.g. overexpression of mTOR or mTORC1/2 or pathogenic variants of tyrosine kinase receptors such as FGFR1 or RET [16–19].

Inhibitors of different components of this signaling cascade, e.g. AZD8055 targeting mTORC1/2 [16], and metformin affecting AKT/mTOR [20] have demonstrated antitumor effects against rodent PPGL cell lines *in vitro*. We previously showed that inhibiting PI3K and mTOR via the combination of alpelisib (BYL719) and everolimus (RAD001), or by using the dual PI3K/mTOR inhibitor (BEZ235), potently suppresses the proliferation of rodent PPGL cell lines *in vitro* [21, 22], and that BEZ235 suppresses the growth of PCCs in the MENX rat model *in vivo* [22]. We and others have extended the evaluation of inhibitors of PI3K (LY294002, BYL719) or of mTOR (AZD8055, everolimus) to patient-derived primary PPGL cultures *in vitro* with promising results [23,24]. However, in two small studies, everolimus showed only modest efficacy in PPGL patients [25,26]. The combination of sunitinib plus the mTOR inhibitor rapamycin showed efficacy in one *SDHB*-mutant mPPGL patient [27].

Dysregulation of cell cycle control is key driver of PPGL tumorigenesis as demonstrated by studies of mice defective in Rb or cyclin-dependent kinase (CDK) inhibitors such as p16, p18, p21, or p27) [28–31]. A germline inactivating mutation in p27 causes the MENX syndrome in rats, characterized by bilateral pseudohypoxic PCCs (100% frequency) and abdominal PGLs (20%) [32,33]. In human primary PPGLs, loss of p27 and cyclin E overexpression have been observed [34,35]. Similarly, loss of *CDKN2A* (p16), resulting in increased CDK4/6 activity and cell cycle progression, is common in sporadic PPGLs and especially in metastatic cases [36,37]. Noteworthy, recent multiomics studies confirm that cell cycle dysregulation is a key feature of mPPGLs [38,39]. Given these molecular alterations, CDK4/6 inhibitors could represent a potential therapeutic avenue for these tumors but have not been tested so far in clinical settings.

Given that the PI3K pathway and cell cycle progression are heavily involved in PPGL pathogenesis, we considered co-targeting both PI3K and CDK4/6 as potential treatment for PPGL. We selected buparlisib (BKM120), a potent and highly specific oral pan-class I PI3Kα inhibitor, and ribociclib (LEE011), an orally available small molecule, which selectively binds the ATP cleft of CDK4 and CDK6, thereby blocking Rb phosphorylation and, consequently, cell cycle progression.

We report that combined PI3K and CDK4/6 inhibition shows stronger antiproliferative effects than single treatments in *in vitro* PPGL models, including primary patient-derived cultures, and demonstrates synergistic antitumor efficacy in PPGL cell-derived mouse xenografts. Transcriptome analysis revealed that the combination treatment uniquely downregulates Foxm1 and mitotic spindle-associated genes, leading to mitotic catastrophe. These genes are highly expressed in human mPPGLs (*versus* non-metastatic cases), and associate with increased metastatic risk regardless of tumor cluster. Thus, dual PI3K-CDK4/6 inhibition holds promise for treating mPPGLs, currently lacking effective therapies.

## Materials and Methods

### Cell culture and treatments

PC12 cells were obtained from ATCC, while MPC cells were provided by Prof. A. Tischler (Tufts University School of Medicine). Cells were maintained in culture as described in Powers, J. et al [40]. Cells in culture were regularly ensured to be mycoplasma-free. Drug treatments were conducted as outlined in figure legends and Suppl. Table 1.

### Drugs

Buparlisib (PI3Ki) (MedChemExpress) and ribociclib (CDK4/6i) (MedChemExpress) were dissolved in DMSO for *in vitro* applications and in a mixture of 1/9 v/v 1-methyl-2-pyrrolidone/ PEG300.

### Human samples

This study was approved by the local ethics committee of the University Hospital of the Technical University Munich (TUM) (Nr. 2024-30-S-SB), the Ludwig Maximilian University (LMU) Munich (Nr. 379-10), University Hospital Carl Gustav Carus Dresden (multicenter prospective monoamine-producing tumor study [PMT]: Nr. EK 189062010; PROSPHEO [NCT03344016]: Nr. EK 210052017) and by ENS@T (European Network for the Study of Adrenal Tumors). Written informed consent was obtained from each patient prior to participation.

Fresh primary tumor tissues were obtained directly after surgery from 6 PPGL patients at the University Hospital of LMU Munich and numbered consecutively. Details are reported in Suppl. Table 2.

For samples in the CNIO cohort, metastatic disease was defined as the occurrence of distant metastases in anatomical locations where chromaffin tissue is normally absent, as per WHO definition [41]. Aggressive disease was established when there was capsular or adipose tissue invasion, and vascular infiltration or evidence of multiple recurrences, without certainty of metastatic disease, as reported [38].

### Rodent care and *in vivo* treatment

*In vivo* studies were approved by the government of Upper Bavaria, Germany (Az.: 55.2.1.54-2532-39-13). MENX-affected rats and immunocompromised CD-1foxn1-nude mice were maintained in agreement with general husbandry rules approved by the Helmholtz Zentrum München. Six-week-old female mice obtained from Charles River were housed in a specific pathogen-free environment at a temperature of 25°C and relative air humidity between 45 and 50%.

The mice (n=40) were injected subcutaneously with 1×10⁶ PC12 cells mixed with Matrigel (1:1 v/v). Twenty days post-injection, mice were randomized into treatment groups (PI3Ki, CDK4/6i, combination, or placebo) and treated daily for 21 days. Tumor volume was measured twice weekly using external calipers. Complete necropsies were performed post-treatment. Collected organs were either fixed in 4% buffered formalin and embedded in paraffin for histological/immunohistological analyses, or stored at –80 °C.

### Primary (human and rodent) 3D organotypic cultures

Tumor tissues (from human patients and MENX rats) were dissociated using the gentleMACS™ Octo Dissociator with heaters using respective tumor dissociation kit (Miltenyi Biotec, Germany) and plated in Ultra-Low Attachment plates. 3D spheroids were allowed to form for 5 days and then treated with drugs for 72 hours.

### Cell proliferation and apoptosis assays

2D Proliferation was measured after 72h of treatment using the CyQUANT® NF Kit (Thermo Fisher Scientific). 3D Viability was assessed at 0, 24, 48, and 72h using the RealTime-Glo™ MT Assay (Promega). Caspase Activity was measured using Caspase-Glo® 9 Assay in 2D and Annexin V Assay in 3D (Promega). 3D spheroid growth was measured by image analysis by ImageJ at days 0, 3, 7, 10 and 14.

### RNA extraction and RealTime quantitative (q) RT-PCR

RNA was isolated using the RNeasy Mini Kit and converted to cDNA using the High-Capacity RNA-to-cDNA Kit. Quantitative RT-PCR was performed using TaqMan inventoried primers and probes (Thermo Fischer Scientific) for the corresponding genes and Fast Advanced Master Mix as previously reported [42]. Used Taqman assays are listed in Suppl. Table 3.

### RNAseq analysis workflow

RNA-seq was performed on PC12 cells treated with PI3Ki (1 μM), CDK4/6i (2 μM), or their combination (at Novogene, Cambridge, UK). Reads were mapped to Rnor_6.0 using the nf-core/rnaseq pipeline. Of three replicates in each group, one sample of the combination group was excluded due to inadequate quality.

RNASeq raw counts were processed as outlined in Suppl. Fig. 1. Differential expression was analyzed using DESeq2 in R. Gene Set Enrichment Analysis (GSEA) was conducted with MSigDB databases. Transcription factor enrichment was analyzed using decoupleR and ChEA3. Protein interaction analysis was conducted using the STRING database.

### Protein extraction and western blotting

Cells were lysed in RIPA buffer with protease and phosphatase inhibitors (Roche Diagnostics) and Western blotting performed as previously described [42] with primary and secondary antibodies and detected using SuperSignal West Pico Chemiluminescent Substrate. (Thermofischer Scientific). Comparative bands were quantified using Image Lab software (Bio-Rad Laboratories). Used antibodies are listed in Suppl. Table 4

### Histochemical staining and Image analysis

Formalin-fixed, paraffin-embedded (FFPE) xenografts were sectioned and stained with an automated immunostainer (Ventana Medical Systems, Tucson, AZ) as described previously [42]. NuSAP immunostaining was analyzed by microscopy, with positive nuclei quantified. Necrotic areas on Hematoxylin and Eosin (H&E) stained tissues were annotated using QuPath software. Immunofluorescence for mitosis visualisation was performed using Alexa 647-tagged secondary antibodies and DAPI. Images were captured at 60x using Olympus FV1200 Confocal microscope.

### Statistical analysis

Study endpoints were represented by bar graphs as mean ± SEM. One-way ANOVA with F-test was used for comparisons among groups. Imaging data were analyzed using paired t-tests. Software used was GraphPad Prism 7-10.

### Limitations of study

Variability in 3D spheroid formation may affect reproducibility. Tumor growth kinetics *in vivo* and experiments concerning primary cultures may vary due to biological heterogeneity. Reagents, equipments and softwares used in the study and respective catalogue numbers are listed in Suppl. Table 5.

## Results

### Combined PI3K and CDK4/6 inhibition suppresses proliferation, migration and invasion and induces apoptosis of PPGL cell lines and primary cells

To identify novel treatments for PPGLs, we targeted two dysregulated processes: the PI3K pathway and cell cycle progression. We tested the sensitivity of PPGL cells to the oral pan-class I PI3K inhibitor buparlisib (PI3Ki) and the selective CDK4/6 inhibitor ribociclib (CDK4/6i) using two cell lines: PC12 (derived from a rat PPGL) [43] and MPC (from a heterozygous Nf1-deficient mouse PPGL) [40]. Both cell lines can serve as aggressive and metastatic models, as MPC cells metastasize in orthotopic or tail vein injection models while PC12 cells can even metastasize from subcutaneous implanted tumors [44].

Cells were treated with clinically relevant doses of PI3Ki, CDK4/6i, or their combination, with DMSO as a control. The drug combination showed the strongest inhibition of proliferation in both cell lines. In PC12 cells, the combination had an IC50 of 0.2 µM at 72 hours, outperforming PI3Ki (IC50: 0.7 µM) and CDK4/6i (IC50: 1.6 µM) (Fig. 1A and Suppl. Fig. 2A; Suppl. Table 2). Similarly, in MPC cells, the combination was most effective (IC50: 0.39 µM) compared to PI3Ki alone (IC50: 0.63 µM) and CDK4/6i alone (IC50: 3.68 µM) (Fig. 1B and Suppl. Fig 2B; Suppl. Table 1). While both drugs individually suppressed cell proliferation, the combination treatment was significantly more effective.

**Figure 1.**
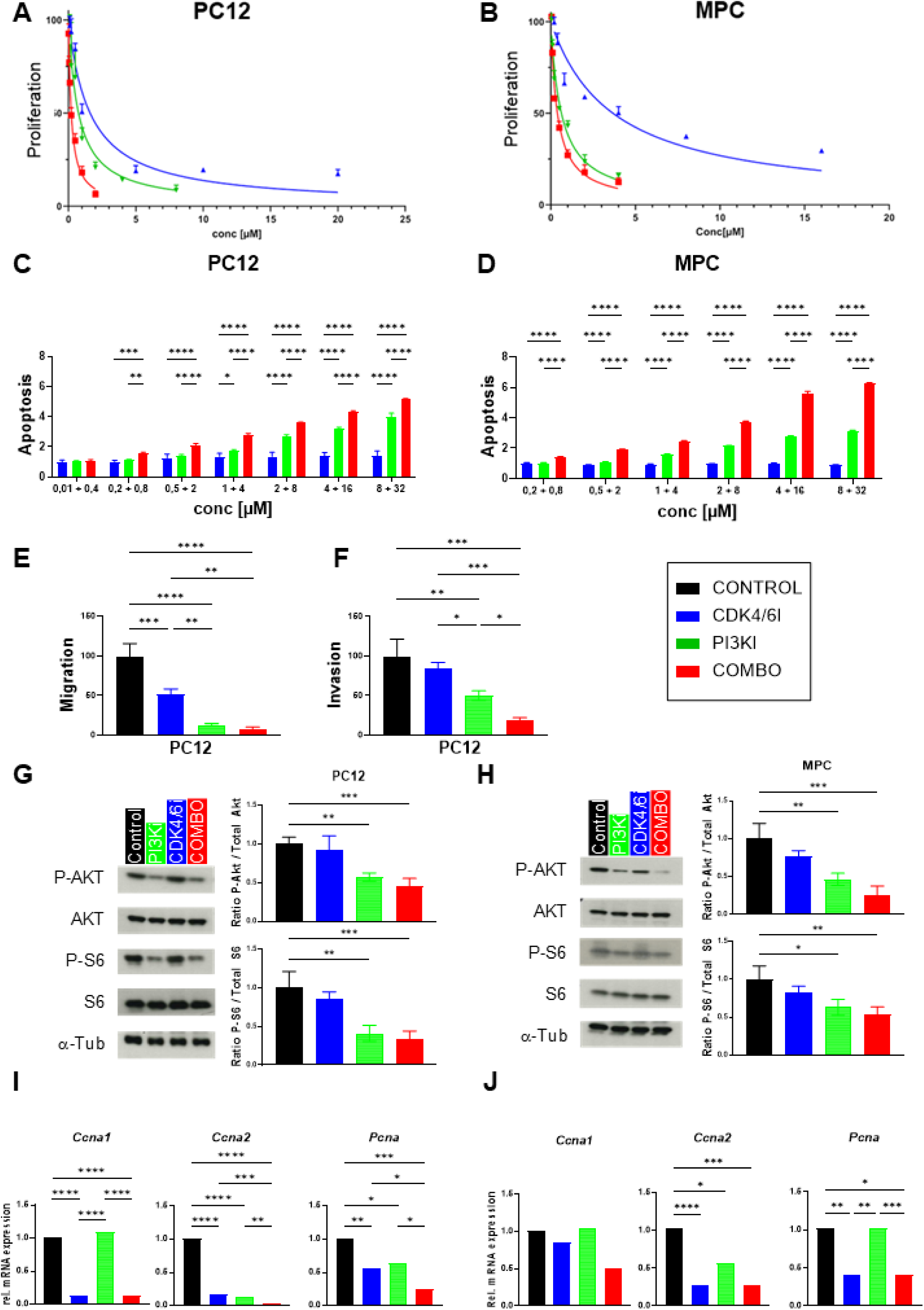
Effect of CDK4/6i and PI3Ki treatment on proliferation, oncogenic features and downstream targets of PC12 and MPC cells grown as 2D cultures. **(A,B)** IC_50_ curves of PC12 (**A**) or MPC (**B**) cells treated with CDK4/6i, PI3Ki, their combination or DMSO control. Cell proliferation was measured after 72h of treatment. For drug concentrations see Suppl. Table S1. The DMSO control was set to 100% and nonlinear regression was used to calculate the IC_50_. Data shown are the mean±SD from 3 independent experiments with 3 technical replicates each. (**C,D**) PC12 (**C**) and MPC (**D**) cells were treated using the indicated concentrations and Caspase 9 activity was measured 72h later. Shown is the relative apoptosis normalized to the DMSO control. Data shown is the mean ± SD from 3 independent experiments with 3 technical replicates each. Statistics: 2way ANOVA. (**E,F**) PC12 cells were plated in chambers without Matrigel to assess migration (**E**), or containing Matrigel to assess invasion (**F**). After 72h of treatment with IC_50_ values of the drugs, chambers were collected, stained and cells counted (3 technical replicates for each condition). Data shown are the mean±SD from 3 independent experiments. Statistics: 1way ANOVA. ns, not significant; *, p< 0.05; **, p< 0.01; ***, p<0.001; ****, p< 0.0001. (**G,H**) Expression and quantification of total Akt, phospho-Akt (pAKT), total S6, phosphor-S6 and α-Tubulin (loading control) in PC12 (**G**) or MPC (**H**) cells treated with the indicated drugs for 72h. Shown is one representative immunoblot out of 3 independent experiments. Band quantification was conducted using Image J. Statistics: 1way ANOVA. ns, not significant; *, p< 0.05; **, p< 0.01; ***, p<0.001; ****, p< 0.0001. (**I,J**) Realtime qRT-PCR for *Ccna* and *Pcna* was conducted on PC12 (**I**) and MPC (**J**) cells with the various drugs/drug combination. RNA was extracted 72h post-treatment. Values were normalized against the DMSO control arbitrarily set to 1. Data are shown as the mean of three experiments ± SD. Statistics: 1way ANOVA. ns, not significant; *, p< 0.05; **, p< 0.01; ***, p<0.001; ****, p< 0.0001.

To evaluate whether the anti-proliferative effects of PI3Ki and/or CDK4/6i in PPGL cells involve increased cell death, we measured Caspase 9 activity in PC12 and MPC cells 72 hours post-treatment. Both PI3Ki alone and the drug combination induced apoptosis in a dose-dependent manner, while CDK4/6i modestly triggered apoptosis only at higher concentrations (Fig. 1C,D and Suppl. Fig 3). Notably, the combination treatment induced apoptosis even at low doses (PI3Ki 0.5 µM + CDK4/6i 2 µM for PC12 and PI3Ki 0.2 µM + CDK4/6i 0.8 µM for MPC), reflecting the combination’s superior efficacy in reducing cell proliferation.

We next assessed the impact of PI3Ki and/or CDK4/6i treatment on clinically relevant oncogenic behaviors, such as migration and invasion, in PPGL cells using the Boyden chamber assay. All treatments significantly reduced PC12 cell migration at 72 hours compared to the DMSO-treated control (Fig. 1E). Notably, PI3Ki alone strongly suppressed migration, while the combination treatment nearly abolished it (Fig. 1E).

We have shown that our treatment approach is able to reduce the proliferation and induce the death by apoptosis of both PC12 and MPC cell lines. To verify that the observed phenotypes were indeed caused by pathway inhibition and not by unspecific effects, we assessed different downstream targets of PI3K/AKT or CDK4/6 pathways. For the former, we analyzed the effect of the drugs/drug combination on the phosphorylation of AKT (at the S473 residue) and S6 (at residue S240/244). Treatment with PI3Ki alone and with the drug combination significantly reduced both P-AKT/total AKT and P-S6/total S6 signal ratios in both cell lines (Fig. 1G,H), confirming PI3K inhibition while. treatment with DMSO (vehicle control) and CDK4/6i had no effect on AKT nor on S6 phosphorylation (Fig. 1G,H).

To validate CDK4/6 inhibition, we analyzed the expression of three genes that are targets of E2F factors and thus should be repressed by CDK4/6 inhibition [45,46], namely *Ccna1* (cyclin A1)*, Ccna2* (cyclin A2), and *Pcna*, (Proliferating Cell Nuclear Antigen). In general, these cell cycle-dependent genes were downregulated more strongly by CDK4/6i and the combination treatment than by PI3Ki alone, as expected (Fig 1 I,J). Only in PC12 cells there was a slightly stronger effect on the expression of *Ccna2* and *Pcna* in the combo group compared to CDK4/6i alone (Fig 1I). Altogether, the drugs inhibited the known direct targets as anticipated, while the higher efficacy of the combination could not be explained by the inhibition of these pathways alone.

### Combined PI3K and CDK4/6 inhibition suppresses the viability and growth of 3D organotypic cultures of PPGL cells

As 3D organotypic culture systems have a higher predictive value due to physiological cell-cell contacts, [47], we extended our treatments to spheroids of PC12 and MPC cells. Cells were treated with different doses of the single drugs or drug combinations or left untreated (DMSO control), and cell viability was assessed 24h, 48h and 72h later. For PC12 cell spheroids, all conditions reduced cell viability *versus* DMSO control starting from the dose 0.2µM PI3Ki and 0.8µM CDK4/6i (Fig. 2A and Suppl. Fig. 4A). The combination was significantly more effective compared to the single drugs for all the tested concentrations. MPC cells spheroids were more sensitive to CDK4/6i than PC12 cells, and the drug combination was effective starting at concentrations of 1µM PI3Ki and 4µM CDK4/6i (Fig. 2B and Suppl. Fig.4B). We also investigated the growth of the 3D organotypic cultures by measuring spheroid sizes at day 0, 3 and 14 days post-treatment (Fig. 2C,D and Suppl. Fig. 5). For MPC cells there was additive effect of the drug combination at inhibiting spheroid size with respect to PI3Ki alone, whereas for PC12 this effect was not observed in terms of spheroid size (Fig. 2C,D and Suppl. Fig. 5).

**Figure 2.**
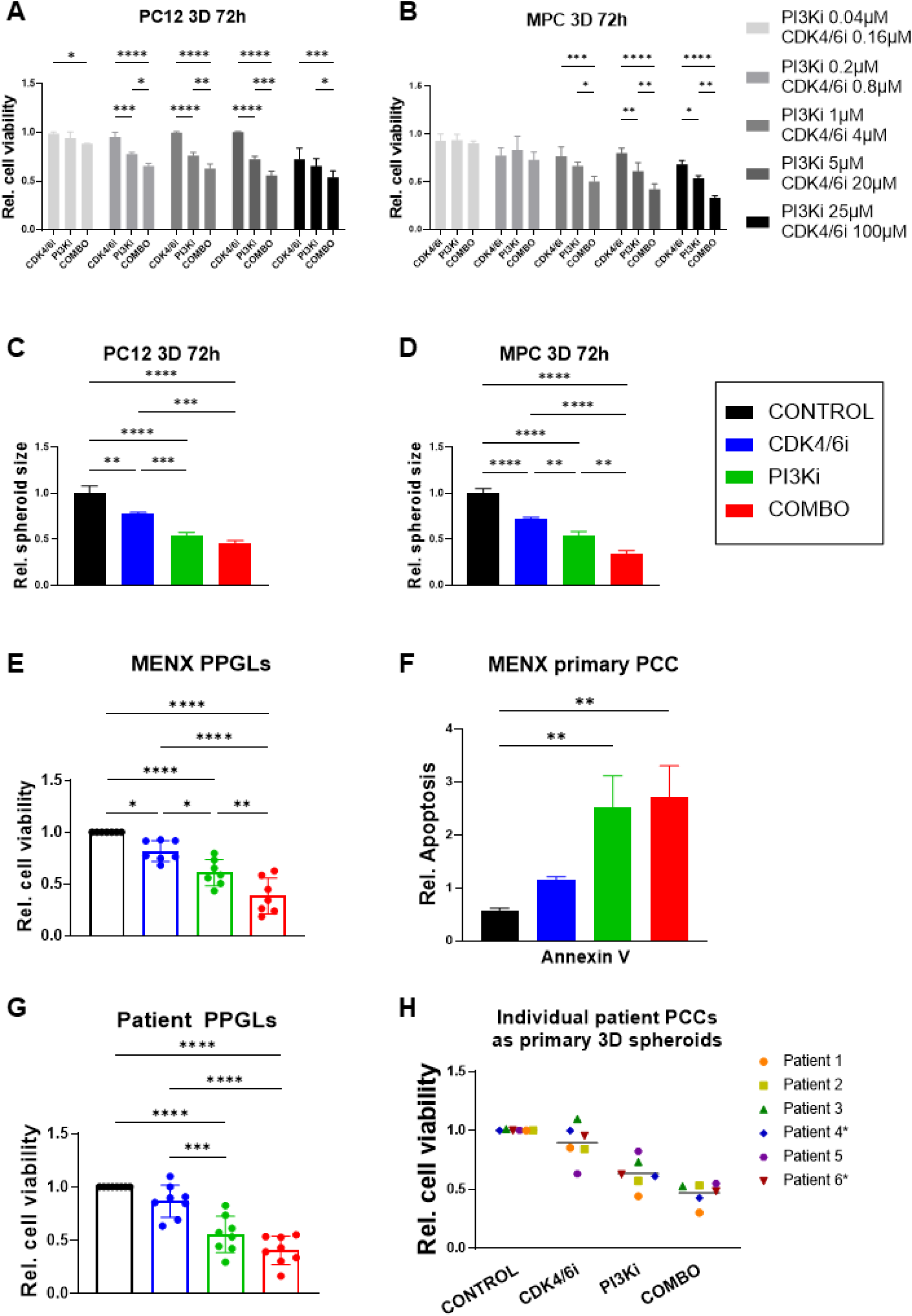
Effect of CDK4/6i and PI3Ki treatment on the viability and invasion of 3D organotypic cultures of PPGL cell lines and rat or human primary tumor cells. **(A, B)** Following spheroid formation, PC12 (**A**) or MPC (**B**) cells were treated with different doses of CDK4/6i, PI3Ki, their combination or DMSO control. Cell viability was assessed 72h later. Reported is the relative cell viability normalized against untreated control (arbitrarily set to 1). Data shown are the mean±SD from 3 independent experiments with 8 technical replicates each. Statistics: 1way ANOVA. ns, not significant; *, p< 0.05; **, p< 0.01; ***, p<0.001; ****, p< 0.0001.(**C,D**) 3D spheroids of PC12 (**C**) and MPC (**D**) cells were treated with the indicated drugs at the concentration of CDK4/6i: 8 μM; PI3Ki: 2 μM, combination: 8μM CDK4/6i + 2 μM PI3Ki. Spheroid size was measured over time and the values at the 3 Day time point are shown. Data shown are the mean ±SD from 3 independent experiments with 8 technical replicates each. Statistics: 1way ANOVA. ns, not significant; *, p< 0.05; **, p< 0.01; ***, p<0.001; ****, p< 0.0001. (**E,F)** Primary cells were isolated from fresh MENX-associated PCC fragments (n=7). 3D spheroids were generated and treated with the indicated drugs at the concentrations reported in **C**. (**E**) Cell viability was measured 72h later and normalized against the initial measurement and the DMSO control. Data shown is the mean ±SD with 5-15 technical replicates each tumor (depending on total amount of cells). Statistics: 1way ANOVA. ns, not significant; *, p< 0.05; **, p< 0.01; ***, p<0.001; ****, p< 0.0001. (**F**) Primary cells were isolated from 3 rats, processed and treated as in **E**. Annexin V signal was measured 30h post-treatment to assess apoptosis. Shown is relative apoptosis normalized to the DMSO control. Data shows the mean ±SD from primary cells of 3 rats with 8 technical replicates each. Statistics: 1way ANOVA. ns, not significant; *, p< 0.05; **, p< 0.01; ***, p<0.001; ****, p< 0.0001.(**G,H**) Primary cells were isolated from 8 fresh patient-derived PPGL and 3D organotypic cultures generated. Spheroids were treated with CDK4/6i (8μM), PI3Ki (2μM), their combination (CDK4/6i 8μM + PI3Ki 2 μM), or DMSO. Cell viability was measured 72h later. Shown is the relative cell viability normalized to the initial measurement and the DMSO control. In (**G**) the mean ±SD from primary cells of 8 patients with 4-8 technical replicates each (depending on total amount of cells) is shown. Statistics: 1way ANOVA. ns, not significant; *, p< 0.05; **, p< 0.01; ***, p<0.001; ****, p< 0.0001. In (**H**) the response of 6 individual patients is illustrated as the mean value.

We assessed drug sensitivity in primary PCC cells from MENX rats, a representative model where homozygous mutants spontaneously develop bilateral PCCs by 7-8 months [32, 48].

We treated 3D primaryPCC spheroid cultures from seven rats with PI3Ki, CDK4/6i, their combination, or DMSO. Due to limited material, a single concentration of each drug or combinationwas used. PI3Ki reduced cell viability by nearly 50%, showing greater efficacy than CDK4/6i, while the combination caused the strongest suppression (Fig. 2E). Apoptosis assays revealed that PI3Ki and the combination therapy induced apoptosis, whereas CDK4/6i alone did not (Fig. 2F).

For an easier clinical translation of our findings, we performed drug testing also on primary patient-derived (ppd) PPGL cultures. We recently showed that ppd PPGL cultures predict the responsiveness of primary tumors to specific drugs, thereby representing a highly valuable translational tool [24]. Six fresh tumor samples, two of them derived from mPPGL patients, were processed to obtain 3D organotypic cultures. The detailed characteristics of each patient are reported in Suppl. Table 2. After spheroid formation, cells were treated with the single drugs, their combination or DMSO control and cell viability was assessed 72h later. Due to limited material, we only used one concentration for each individual drug and for the combination. While CDK4/6i showed a positive trend (albeit not-significant) in reducing cell viability, both PI3Ki and the drug combination significantly and strongly reduced patient-derived primary PPGL cell viability *versus* DMSO-treated control cells (P<0.001) (Fig. 2G,H). There was a trend for the combination to be more effective than PI3Ki alone, but it wasn’t statistically significant. Noteworthy, cells derived from the two mPPGLs (indicated with an asterisk) responded well to this combination therapy (Fig. 2H).

Altogether, these results attest to the efficacy of targeting both PI3K and CDK4/6 in suppressing the viability of primary rat and human PPGL 3D organotypic cultures.

### Does the efficacy of the PI3Ki-CDK4/6i combination extend to other drugs belonging to these classes?

We demonstrated that the combination of a pan-class I PI3Ki (buparlisib) with a CDK4/6i is highly effective against PPGL *in vitro* including human primary mPPGL cells. Class I PI3Ks consist of a regulatory subunit bound to a p110 catalytic subunit (p110α, β, γ or δ). This binding maintains the p110 subunits in an inactive form until their regulatory subunits are activated by upstream signaling [49]. In addition to pan-PI3Ki, also isoform-specific PI3Ki have been developed and several are used in clinical practice, e.g. alpelisib specifically targeting PI3Kα, or duvelisib inhibiting PI3Kδ. We wondered whether the anti-proliferative effects of inhibiting both PI3K and CDK4/6 pathways in PPGL cells could be extended to isoform-specific PI3Ki. Thus, we treated PC12 cells with alpelisib and duvelisib alone or in combination with ribociclib. The results showed that the efficacy of duvelisib, but not of alpelisib, at inhibiting PPGL cells proliferation is enhanced when combined with ribociclib (Suppl. Fig. 6A).

In complementary experiments, we evaluated two alternative CDK4/6 inhibitors, e.g. palbociclib and abemaciclib, in combination with buparlisib, but could not reproduce the significant increase in the efficacy of the drug combination *versus* PI3Ki alone that we observed using ribociclib (Suppl. Fig. 6B). Due to the limited material available from human PPGL tumors, we could not extend these experiments to primary human 3D organotropic cultures to examine, whether this dependency on ribociclib is PC12 cell line specific or might be a consequence of a specific drug effect only with this compound.

### Combined PI3K and CDK4/CDK6 shows strong antitumor activity *in vivo*

The data presented above indicates that combined PI3K and CDK4/CDK6 inhibition is effective against various representative *in vitro* models of PPGL. Next, we tested whether the combination of PI3Ki and CDK4/6i is also effective *in vivo* in a PC12 cell-derived xenograft model. PC12 cells were implanted in the flanks of CD1foxn1-nude immunodeficient mice (n=40 in total). When the tumors reached approximately 100mm^3^ in volume, mice were randomized in 5 different groups (n=8 mice/group). Tumor-bearing mice were treated with PI3Ki alone (25mg/Kg bw), with CDK4/6i alone (75mg/Kg bw), with both drugs at the same doses used for the single treatments (“regular dose” regimen), with both drugs at reduced dose (i.e. PI3Ki =12.5mg/Kg bw; CDK4/6i=25 mg/Kg bw – “lower dose” regimen) or with vehicle control (1 volume of 1-methyl-2-pyrrolidone and 9 volumes of PEG300) for 21 days. Tumor volume was longitudinally measured during treatment by external caliper. At the end of the treatment, tumorgrafts were collected, fixed and FFPE tissue blocks were generated for *ex vivo* analyses. Treatment with CDK4/6i or PI3Ki as single agents suppressed tumor growth, leading to a reduction in tumor volume of –18% and –39%, respectively, with volume in the latter group being statistically significantly smaller than in the control group (P<0.001) (Fig. 3A). Remarkably, administration of both drugs at the regular dose resulted in a rapid tumor regression: up to 78% tumor volume shrinkage *versus* vehicle (P<0.001) (Fig. 3A). More importantly, the higher therapeutic efficacy of the drug combination *versus* single drugs was also achieved by administering half of the dose of PI3Ki, together with a third of the dose of CDK4/6i, i.e. the lower dose regimen (Fig. 3A). This indicated that combined targeting of PI3K and CDK4/6 has a pronounced synergistic effect on the inhibition of PC12 cell-derived xenografts’ growth *in vivo*.

**Figure 3.**
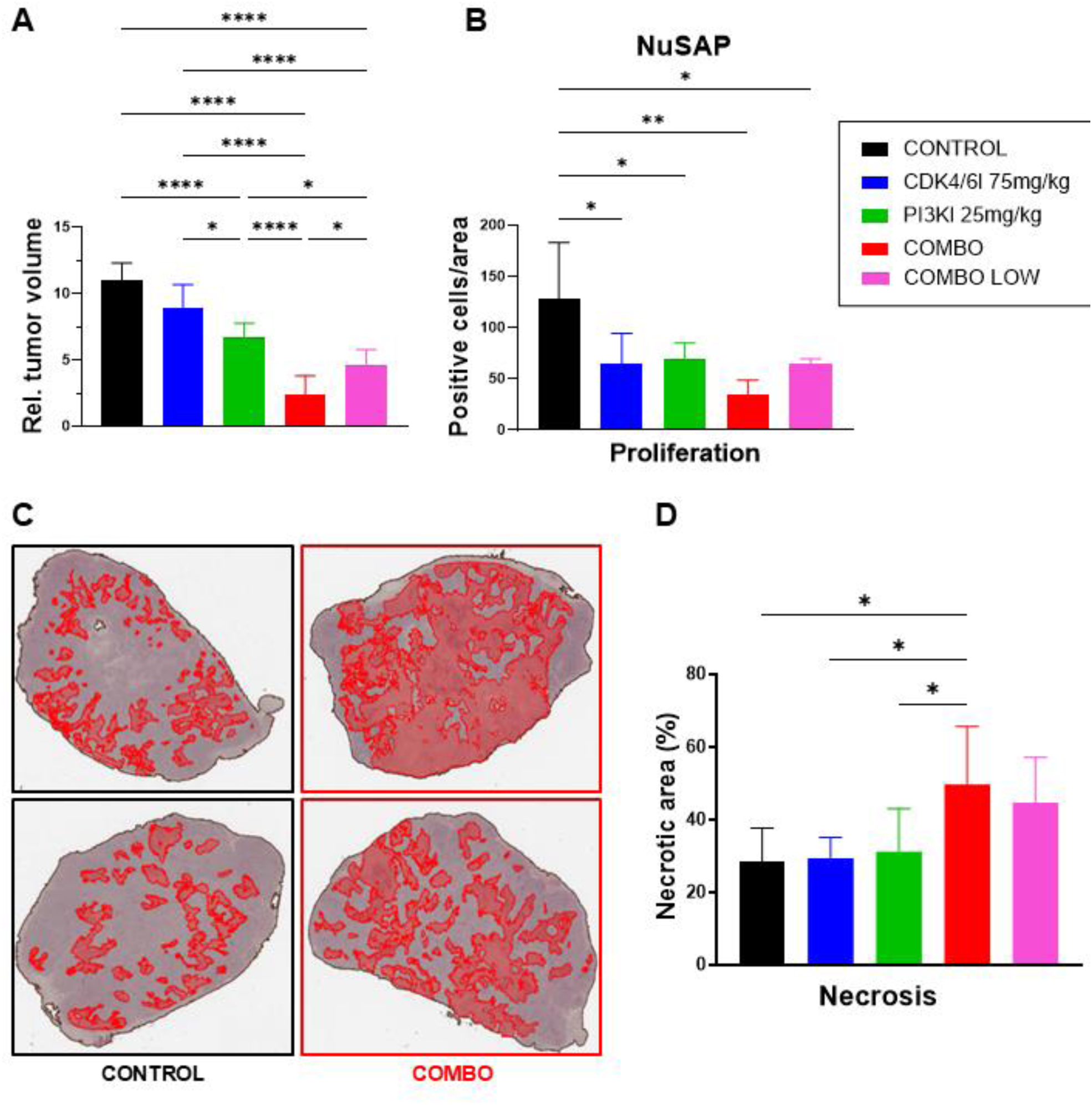
Effect of CDK4/6i and PI3Ki treatment on the growth of PC12-derived mouse xenografts. (**A**) PC12 cells were injected in CD-1*^foxnu^* female mice. When tumors reached the size of mm^3^, mice were randomized into the 5 treatment groups, color-coded as in B. Drugs were administered by oral gavage daily for three weeks at the following concentrations: CDK4/6i 75mg/Kg; PI3Ki 25mg/Kg; combination “regular dose”: CDK4/6i 75mg/Kg + PI3Ki 25mg/Kg; combination “low dose”: CDK4/6i 25mg/Kg + PI3Ki 12.5mg/Kg. Tumor volume was measured 2X/week by external caliper. Relative tumor volume compared to the beginning of treatment for mice in each treatment group (n=8/group). Shown is the mean ±SD from tumors of 6-8 mice per group after outlier removal in GraphPad Prism using ROUT method. Statistics:1way ANOVA. ns, not significant; *, p< 0.05; **, p< 0.01; ***, p<0.001; ****, p< 0.0001. (**B**) FFPE sections of 4 independent tumorgafts per treatment group were stained with the anti-NuSAP antibody to detect dividing cells. Upon immunofluorescence (20X magnification), the number of positive cells per 10fields of view were counted. Shown is the mean ± SD. Statistics: 1way ANOVA. ns, not significant; *, p< 0.05; **, p< 0.01; ***, p<0.001; ****, p< 0.0001. (**C,D**) Automated assessment of necrotic areas in the tumorgrafts. (**C**) Sections of. independent tumorgrafts per treatment group were stained with H&E and scanned with a software to detect necrotic areas. In the exemplary images, necrotic areas are in red. (**D**) Quantification of the necrotic areas in the 5 treatment groups, color-coded as in **B**. Statistics: 1way ANOVA. ns, not significant; *, p< 0.05; **, p< 0.01; ***, p<0.001; ****, p< 0.0001.

**Figure 4.**
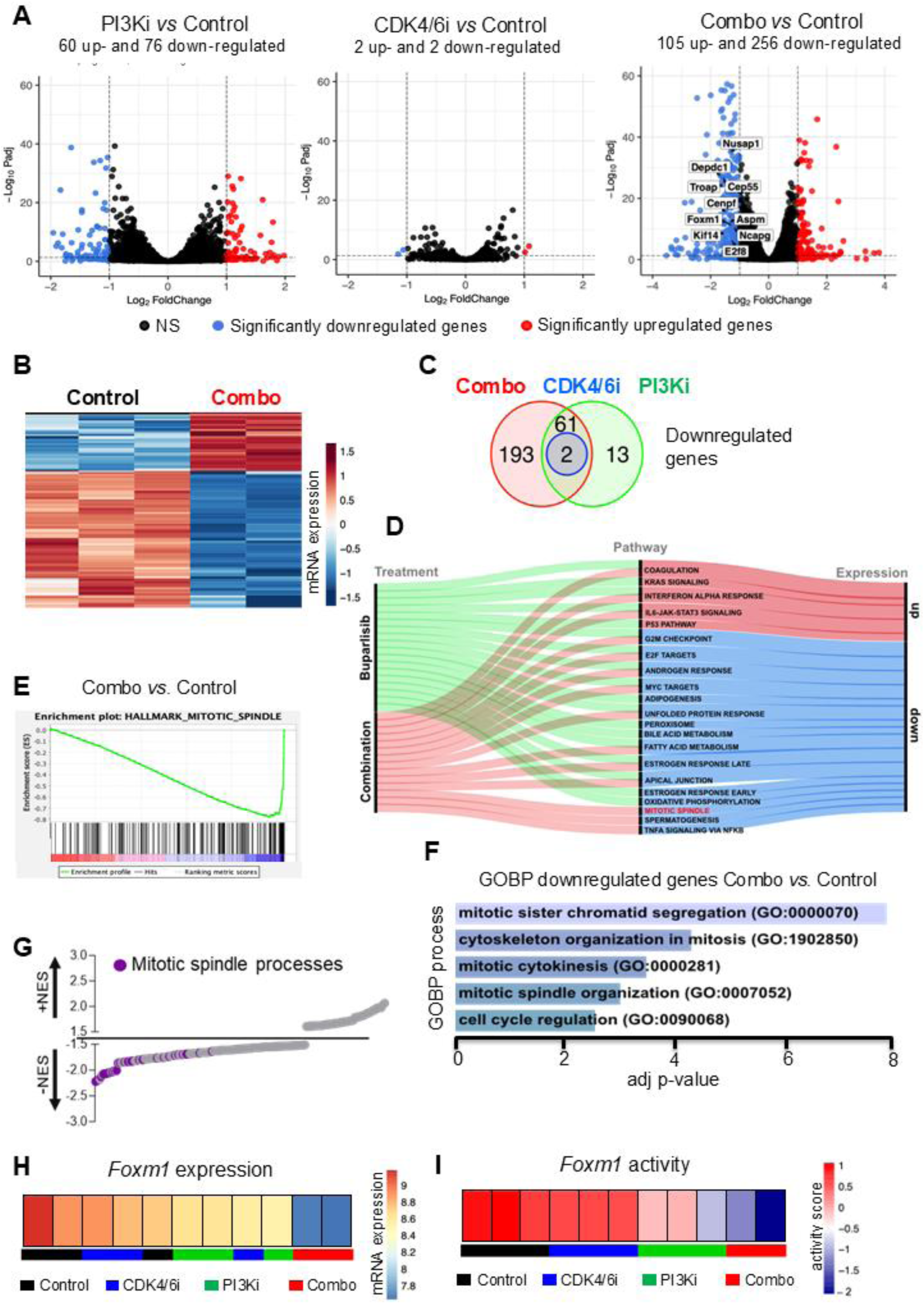
Transcriptomic analyses of PPGL cells treated with the drugs/drug combination. (**A**) Volcano plots depicting the DGEs for the comparison between PI3Ki-, CDK4/6i-, and combination-treated PC12 cells *versus* untreated control samples. The number of DGEs is indicated. (**B**) Heatmap showing hierarchal clustering of the DGEs between combination and control samples. (**C**) Venn diagram illustrating the intersection of the downregulated genes in the three treatment groups when compared with control samples. (**D**) Alluvial plot showing the overlap of significantly enriched (p< 0.05; FDR<0.25) upregulated (red) and downregulated (blue) pathways between the PI3Ki and combination groups. The mitotic spindle pathway, enriched exclusively in the combination, is highlighted in red. (**E**) GSEA enrichment plot for the only category exclusively downregulated in combination *versus* control from Hallmark. (**F**) Bar plot showing the top five enriched processes of downregulated genes unique to the combination on GOBP. In parenthesis is the associated GO term code; below the adjusted p-value is shown. The analysis was conducted using Enrichr. (**G)** GSEA plot showing the NES of significantly enriched gene sets in the mitotic spindle processes category (p< 0.05; FDR<0.25) from MSigDB. Each dot represents a gene set and in purple are highlighted the gene lists related to mitotic spindle processes. (**H-I**) Heatmaps showing the gradient of *Foxm1* expression (**H**) or Foxm1 activity (**I**) in PC12 cells following the indicated treatments.

To investigate if the suppression of tumor growth observed *in vivo* was associated with the inhibition of tumor cell proliferation, we conducted IHC for the proliferation marker NuSAP on xenograft tissue [50, 51]. The number of positive nuclei was counted in various sections of at least 3 independent tumors from each treatment group and was then compared among the groups. Tumors of treated mice (regardless of the drug regimen) showed a general decrease in the number of NuSAP-positive nuclei *versus* untreated controls (Fig. 3B). The strongest reduction in cell division was observed in mice treated with the regular dose of the combination therapy (Fig. 3B). This data is in line with the kinetic of tumor growth assessed by caliper measurements.

To determine whether the drug treatments also promoted cell death *in vivo*, we quantified the necrotic areas of eight tumorgrafts per group of whole tissue mounts. The combination of PI3Ki and CDK4/6i (regular dose) significantly increased the necrotic areas in the tumors when compared with both control and CDK4/6i-treated samples, but also *versus* PI3Ki alone (Fig. 3C,D). The lower dose regimen promoted greater cell necrosis than the single drugs, though the difference was not statistically significant (Fig. 3C,D). These data suggest that the combination treatment elicited strong antiproliferative and cytotoxic effects on PPGL cells *in vivo.* No weight change or other adverse effect on mice health was noticeable due to the treatment.

### Deregulation of mitotic spindle genes is a hallmark of combined PI3K and CDK4/CDK6 inhibition

We demonstrated that dual PI3K+CDK4/6 inhibition exhibits potent, antitumor effects *in vitro* and *in vivo*. To investigate the underlying molecular mechanisms, we performed RNASeq on PC12 cells treated with buparlisib, ribociclib, their combination, or DMSO for 72 hours at IC_50_ doses. RNASeq analysis (Suppl. Fig. 1) revealed distinct clustering of treatment conditions via PCA, with the combination group differing most from controls, followed by PI3Ki and CDK4/6i (Suppl. Fig. 7A-B). DESeq2 analysis identified the highest number of differentially expressed genes (DEGs) in the combination group *versus* control (n = 361), mostly downregulated (n = 256), compared to fewer DEGs in the PI3Ki group (n = 136) and minimal changes in CDK4/6i (n = 4) (p-adj < 0.05 and |log_2_FoldChange| > 1) (Fig. 4A-B, Suppl. Fig. 7C). These results indicate that the combination treatment exerts a synergistic effect on the transcriptome.

To better understand the distribution of DEGs across treatment groups, we performed an intersection analysis. Considering the downregulated genes, the combination group exhibited the most extensive overlap with PI3Ki (n = 61) (Fig. 4C). Albeit the CDK4/6i group demonstrated minimal impact on gene downregulation, remarkably the combination treatment uniquely downregulated 193 genes (Fig. 4C). For the upregulated genes, the majority were shared between the PI3Ki and combination treatment groups (n = 38), reflecting their similar gene expression profiles (Suppl. Fig. 8A).

In the overall analysis, the combination group had the largest number of unique DEGs (n = 259), followed by the comparison PI3Ki-combination (n = 99) (Suppl. Fig. 8B). The PI3Ki treatment alone contributed 34 unique DEGs, while the intersection of all three treatments revealed only 3 common DEGs. CDK4/6i accounted for just one unique DEG (n = 1) (Suppl. Fig. 8B).

To elucidate the biological processes and pathways affected by the treatments, we conducted enrichment analysis using GSEA and GO. Pathways were deemed significant if they met thresholds of p < 0.05 and FDR < 0.25, ensuring robust interpretation.

Given the few DEGs in the CDK4/6i group, we focused on the PI3Ki and combination groups (*versus* control DMSO-treated cells) for GSEA Hallmark analysis further. Both treatments strongly downregulated key cell cycle pathways, including the G2M checkpoint, E2F targets, and mTORC signaling, consistent with *in vitro* results and supporting the hypothesis that the combination therapy inhibits tumor proliferation by targeting essential cell cycle pathways (Suppl. Fig. 9A,B). Notably, the mitotic spindle pathway was significantly enriched in the combination group, but absent in individual treatments (Fig. 4D,E), suggesting that dual PI3K and CDK4/6 inhibition uniquely disrupts spindle formation, enhancing effects on cell division. GSEA enrichment plots showed a significant negative Enrichment Score (ES) for the mitotic spindle pathway, G2M checkpoint, and mTORC signaling upon dual inhibition (Suppl. Fig. 9C), confirming the impact of the combination treatment (p < 0.05, FDR < 0.25).

No universal signature emerged for the pathways upregulated across all treatments (Suppl. Fig. 9A,B). However, the PI3Ki and combination groups displayed enrichment in immune-related pathways, including IL6-JAK-STAT3 signaling and the interferon-alpha response.

To validate the GSEA findings, we conducted a comprehensive GO analysis of Biological Processes (GOBP) using ClusterProfiler focusing on the combination treatment group, which showed the most pronounced changes in gene expression. Both up– and downregulated genes were considered. The GOBP analysis confirmed key results from GSEA, with strong enrichment in the combination of pathways related to the G2M transition and cell cycle regulation, essential for cell proliferation and tumor suppression (Suppl. Fig. 10). Additionally, the GOBP analysis highlighted processes linked to chromosomal segregation, such as spindle assembly and chromosome alignment during mitosis, in agreement with the mitotic spindle pathway being uniquely enriched in the combination group on GSEA (Fig. 4F,G). Together, these results indicate that the combination treatment not only disrupts the cell cycle but also impairs critical mechanisms of chromosomal integrity.

To explore the biological significance of the genes deregulated by combined PI3K and CDK4/6 inhibition, we performed an enrichment analysis using enrichR considering the 259 protein-coding genes exclusively differentially expressed in the combination group *versus* control (Suppl. Fig. 8B), including 66 upregulated and 193 downregulated genes (Fig. 4C and Suppl. Fig. 8A). The downregulated genes unique to the combination treatment showed significant enrichment in pathways related to the mitotic spindle and cell division, including chromosome segregation, cytokinesis, and spindle organization in GO Cellular Component (GOCC) and GO Molecular Function (GOMF) (Fig. 4F and Suppl. Fig. 11A,B). This confirms the disruption of mitotic processes observed in the GSEA analysis, explaining the antiproliferative activity in PPGL cells. Enrichment analysis of upregulated genes across GOBP, GOCC, and GOMF categories highlighted involvement in vesicular transport, vacuole organization, and autophagy, without a shared molecular signature (Suppl. Fig. 11C-E).

To identify potential upstream regulators of gene expression signatures, we performed a transcription factor (TF) analysis using DecoupleR for PI3Ki and combination groups and ranked these TFs based on their estimated activity levels, inferred from the expression of their downstream target genes.

In the PI3Ki treatment group, several TFs were significantly less active than in the control group, including SREBF1/2 (lipid metabolism), NFYA (cell cycle progression), PPARG (fatty acid metabolism), HIF1A (hypoxia response), and NR1H3 (cholesterol regulation), ATF6 (unfolded protein response), E2F3 (cell cycle), and NFE2L2 (oxidative stress) (Suppl. Fig. 12A). Conversely, active TFs in PI3Ki-treated *versus* control cells included VHL (tumor suppression), IRF1 (immune response), STAT2 (cytokine signaling), and FOXO3 (stress response), alongside others like HAND2 (differentiation) and HOXA1 (embryonic development) (Suppl. Fig. 12A).

In the combination treatment group, several TFs were significantly less active than in the control cells, particularly those involved in cell cycle regulation, including E2F1-5, SREBF1/2, SRSF2 (splicing and growth), MYC (cell growth), FOXM1 (mitosis), TFDP1, NFYA, KCNIP3, and NR1H3 (Suppl. Fig. 12B). Conversely, TFs with higher activity upon dual inhibition (*versus* control) included FOXI1, ZNF331, STAT2, PHOX2B, IRF1, MXI1 (transcriptional repression), MAF (differentiation), ZHX2, ARX, and HAND2 (Suppl. Fig. 12B).

To validate the findings obtained for the combination treatment, we conducted additional analyses using Enrichr with the ChEA2022 and ENCODE databases, as well as with ChEA3 The results indicated that downregulated genes were primarily regulated by FOXM1 and E2F4, resulting in disrupted cell cycle and mitotic processes (Suppl. Fig. 12C,D and 13).

### The Foxm1 interactome is a target of combined PI3K and CDK4/6 inhibition

Among the TFs whose activity was strongly reduced by the combination treatment, Foxm1 attracted our attention given its role in mitotic progression, spindle assembly, and chromosomal segregation [52], the processes most downregulated by this treatment. When assessing both Foxm1 activity and expression across all treatment, we saw that both progressively decline going from control to combination-treated cells (Fig. 4H,I).

We then mapped the interactome of Foxm1 with genes regulating the mitotic spindle and cell division using ClusterProfiler. We identified a set of genes whose expression correlated with *Foxm1* activity across the various drug treatments: *Aspm* (assembly factor for spindle microtubules)*, Cenpf* (centromere protein F)*, Cep55* (centrosomal protein 55), *Depdc1* (DEP domain containing 1)*, E2f8* (E2F transcriptiona factor 8)*, Kif14* (kinesin family member 14)*, Mad2l1* (mitotic arrest deficient 2 like 1)*, Ncapg* (condensin complex subunit G)*, Nusap1* (nucleolar and spindle associated protein 1) and *Troap* (trophinin associated protein) (Fig. 5A,B). These genes are all implicated in regulating the mitotic spindle, chromosomal segregation and cell division, and some are known targets of Foxm1, such as *Cep55*, *Cenpf, Kif14, Ncapg*, *Nusap1*. The interaction between Foxm1 and these 10 genes was also validated using the STRING database (Suppl. Fig. 14A,B). This analysis confirmed that all the selected genes are co-expressed with *Foxm1* (black lines). Although Foxm1 has not been demonstrated to be the primary regulator of each one of these genes, they form a complex with experimentally validated connections (pink lines). (Suppl. Fig. 14A,B).

**Figure 5.**
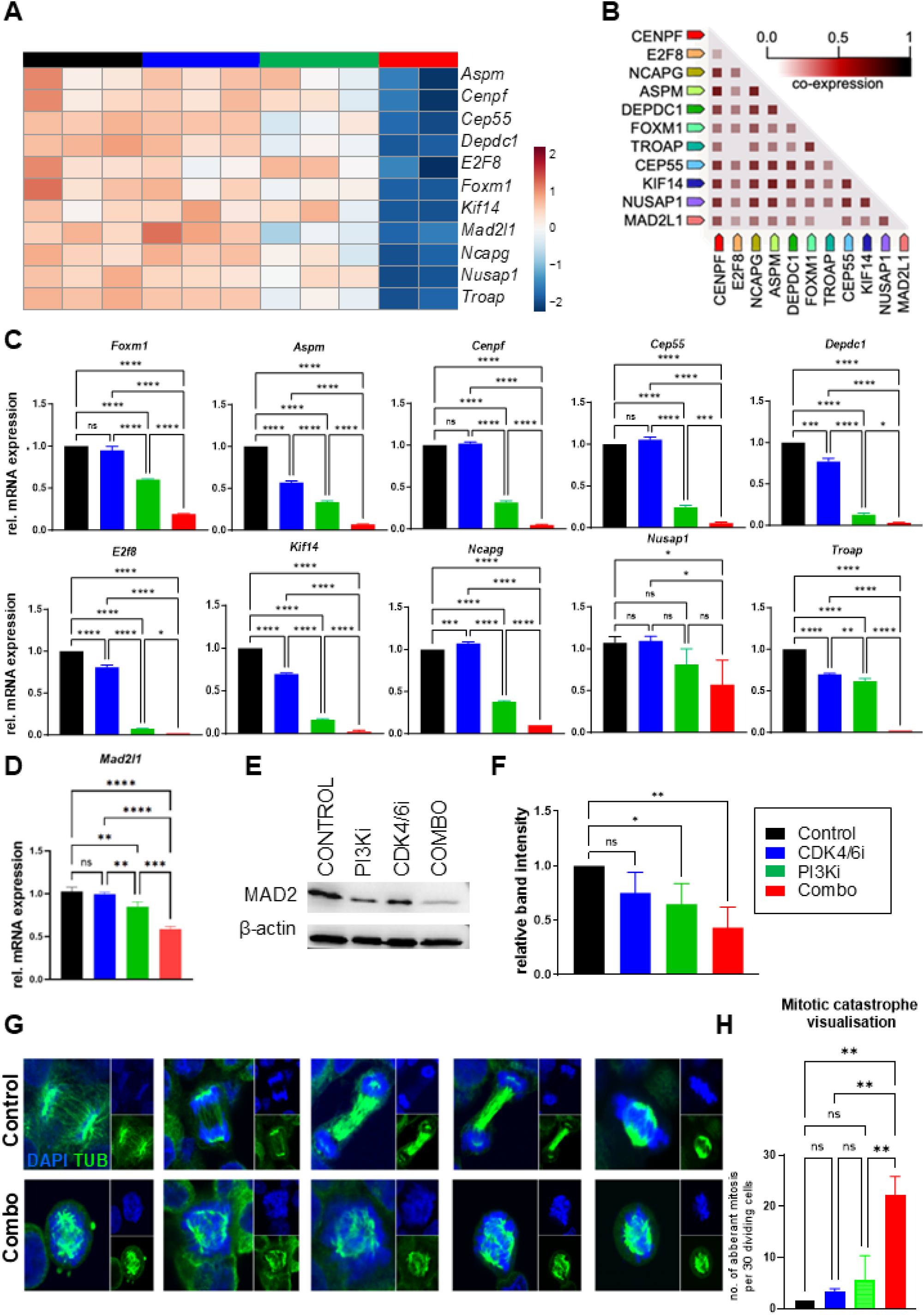
The combination treatment associates with a signature of dysregulated mitotic genes and with mitotic catastrophe in PPGL cells. (**A**) Heatmap showing the expression of selected mitotic spindle-related genes in PC12 cells treated with CDK4/6i, PI3Ki, their combination or with DMSO control based on RNASeq. A gradual decrease in expression can be observed, starting from the untreated (control) samples (black), followed by samples treated with CDK4/6i (blue), PI3Ki (green), and the combination (red). (**B**) Selected genes co-expression plot, generated by STRING, illustrates the relationships among genes based on their co-expression scores calculated on Homo sapiens RNA expression patterns, and on protein co-regulation provided by ProteomeHD. The color scale indicates the degree of co-expression. (**C**) Validation of gene expression of *Foxm1* and its downstream targets in PC12 cells following drug treatments. Genes from the signature shown in A were amplified by realtime qRT-PCR: *Aspm, Cenpf, Cep55, Depdc1, E2F8, Kif14, Ncapg, Nusap1, Troap*. RNA was extracted 72h post-treatment. Values were normalized against the DMSO-treated control arbitrarily set to 1. Shown is the mean±SD. Statistics: 1way ANOVA. Only statistically significant comparisons are indicated with lines and asterisks. (**D**) Expression of *Mad2l1* by realtime QRT-PCR in samples parallel to **C**. (**E**) Western blot analysis of MAD2 protein was performed in PC12 cells treated with 1μM PI3Ki, 4μM CDK4/6i, their combination or left untreated for 72h. (**F**) The relative Mad2l1 protein levels from E were quantified as ratios to beta-actin and normalized to the DMSO-treated control. Results represent the average of three independent experiments; bars, SE. Statistics: one way ANOVA. (**G**) Characterization of mitotic spindle defects in PC12 cells following PI3Ki and CDK4/6i treatment in combination. PC12 cells cultured on Poly-L-lysine coated coverslips were either untreated or treated with 25 mg/Kg PI3Ki and 75 mg/Kg CDK4/6i in combination for 24 h. Cells were then fixed, permeabilized and immunostained with an antibody against α-tubulin (Green). Nuclei were counterstained with 4’,6-diamidino-2-phenylindole (Blue). Mitotic cells were visualized with Leica TCS SP5 (×63 magnification). Exemplary confocal images are shown. (**H**) Quantification of the number of aberrant mitoses in PC12 cells treated as in **G**. At least 30 mitotic cells were captured per condition. Statistics: 1way ANOVA. ns, not significant; *, p< 0.05; **, p< 0.01; ***, p<0.001; ****, p< 0.0001.

The expression of the above identified 11-gene signature was verified by qRT-PCR on independent RNA samples obtained upon treating PC12 cells with the two drugs alone or in combination (Fig. 5C): although their expression could be reduced by PI3Ki alone, the combination always led to a more potent downregulation of the selected genes. Noteworthy, the combination of PI3Ki + CDK4/6i not only reduced the levels of *Mad2l1* mRNA, an important factor regulating mitotic spindle assembly, in PC12 cells more strongly than the individual drugs (Fig. 5C,D), but also led to a stronger reduction of the encoded protein (Fig. 5E,F).

Given that PI3Ki + CDK4/6i suppresses the expression of genes essential for accurate assembly and function of the mitotic spindle, we hypothesized that this treatment regimen might disrupt mitotic fidelity. To address this hypothesis, we stained α-tubulin in PC12 cells previously synchronized by starvation, released and then treated with DMSO, PI3Ki and CDK4/6i alone or in combination. We observed that PPGL cells treated with the combination therapy, but not with the single agents, display a significant increase of aberrant mitotic figures (Fig. 5G,H and Suppl. Fig. 15), which resemble previously reported features of various human cancer cells following loss or repression of FOXM1 [52, 53]. Thus, the inhibition of both PI3K and CDK4/6 pathways promotes mitotic catastrophe *via* the downregulation of transcriptional networks critical for cell division, and this likely explains the higher efficacy of the combination therapy *versus* the single drugs.

Altogether, our molecular analyses have identified molecular footprints predictive of the response of PPGL to combined PI3K and CDK4/6 inhibition.

### Predictive genetic signatures in human metastatic (m)PPGL

We noticed that *Foxm1* and *Troap*, genes of the identified signature, appeared in published transcriptomic datasets comparing metastatic and not metastatic PPGLs [39, 54, 55]. Cancer cells especially with chromosomal instability need to override the mitotic cell cycle checkpoints to ensure rapid proliferation under genetic stress. Therefore, we speculated, that metastatic tumors might rely specifically on the functionality of the mitotic spindle pathway proteins, which then represent an Achilles heel of aggressive tumors. To verify whether the signature mitotic spindle gene list is important in PPGLs we re-analyzed the transcriptome of our large CNIO cohort, including 52 metastatic, 7 clinically aggressive, and 39 non-metastatic cases. We identified 553 DEGs (|log2FoldChange|>1; p-adj < 0.05), with 216 downregulated and 337 upregulated in metastatic samples (Suppl. Fig. 16A). Unsupervised clustering and GSEA Hallmark analysis revealed positive enrichment of genes related to the G2M checkpoint, E2F targets, mitotic spindle, and mTORC signaling in mPPGLs, correlating with their aggressive behavior (Fig. 6A, Suppl. Fig. 16B). In contrast, genes involved in DNA repair, KRAS signaling, and interferon response showed negative NES in metastatic samples (Fig. 6A).

**Figure 6.**
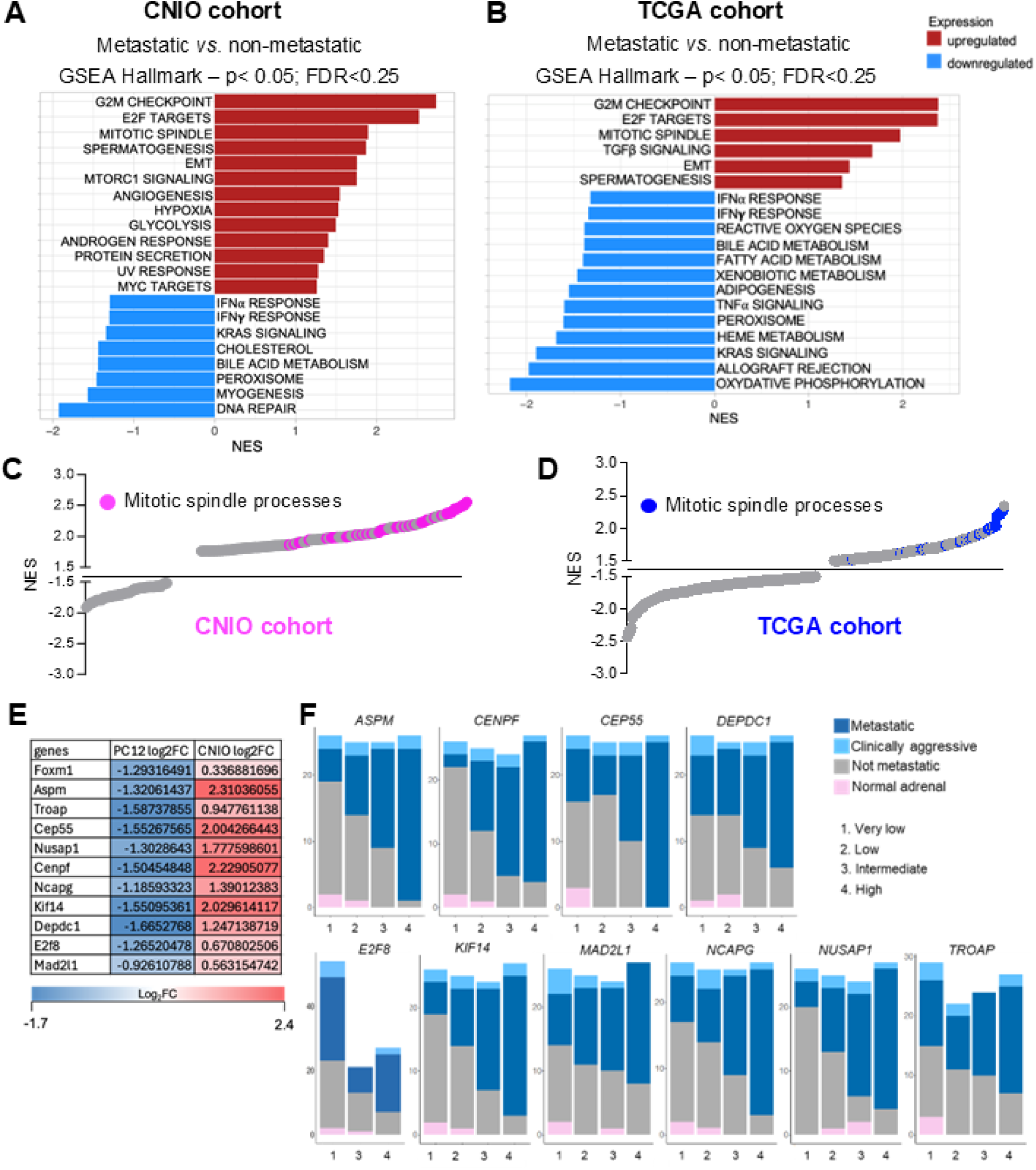
Upregulation of mitosis-related genes in human metastatic *versus* non-metastatic PPGLs. (**A-B**) Bar plots showing the NES (p<0.05; FDR<0.25) on GSEA Hallmarks in metastatic *vs.* non-metastatic patients from the CNIO cohort (**A**) or the TCGA cohort (**B**) on GSEA Hallmark. (**C-D**) Dot plot showing the NES of significantly enriched gene sets in metastatic *vs.* non-metastatic patients from the CNIO cohort (**C**) or the TCGA cohort (**D**) relative to mitotic spindle processes (p<0.05; FDR < 0.05) in GOBP from C5 repository of MSigDB. Each dot represents a gene set and in pink (**C**) or blue (**D**) are highlighted the gene lists related to mitotic spindle processes. (**E-F**) Log2FC expression table (**E**) comparing the expression of mitosis-related genes in combination (CDK4/6i+PI3Ki) treated PC12 cells (relative to control) and metastatic PPGLs from the CNIO cohort (relative to non-metastatic cases). Stacked bar plot (**F)** summarizing the expression levels of the selected gene list across different tumor behaviors in PPGL TCGA dataset.

Remarkably, similar results were obtained applying the GSEA Hallmark analysis to the 16 aggressive/metastatic and the 142 non-metastatic samples belonging to the TCGA PCPG cohort: G2M checkpoint and mitotic spindle were the most dysregulated processes (|log_2_FoldChange|> 1 and a p-adj<0.05) (Fig. 6B and Suppl. Fig. 17A).

Further, GOBP analysis from MSigDB corroborated these findings by revealing significant enrichment in genes related to sister chromatid separation, metaphase-to-anaphase transition, cell cycle checkpoint organization, and various catabolic processes in metastatic *versus* non-metastatic PPGLs in both patients’ cohorts (Suppl. Fig. 16C and 17B). The positive NES of the mitotic spindle pathway in mPPGL in both human tumor cohorts is illustrated in Fig. 6C,D. Interestingly, the mitotic spindle category showed an opposite enrichment in human metastatic *versus* non-metastatic PPGLs when compared with combination-treated PC12 cells (*versus* control), as in the latter it was among the most downregulated processes (see Fig. 4).

Next, we conducted TF analysis on the CNIO cohort samples (containing more metastatic samples than the TCGA cohort) to evaluate the activity of key upstream regulators. Although *FOXM1* expression was not significantly different between the patient groups (log_2_FoldChange= 0.34), its activity, which is mainly regulated by post-transcriptional phosphorylation [56], was markedly elevated in metastatic samples (Suppl. Fig. 18). Other TFs, such as E2F1-5, HCFC1, and REST also showed increased activity in mPPGL, likely contributing to enhanced cell cycle progression markers typical of these aggressive tumors (Suppl. Fig. 18).

We then had a closer look to the human homologues of the mitosis-related genes found downregulated in PC12 treated with combined PI3Ki + CDK4/6i, i.e. *ASPM, CENPF, CEP55*, *DEPDC1*, *E2F8*, *FOXM1*, *KIF14*, *MAD2L1, NCAPG, NUSAP1*, *TROAP*. In the CNIO cohort, these genes were generally more highly expressed in metastatic than in non-metastatic samples by unsupervised clustering. *ASPM*, *CENPF, CEP55*, *DEPDC1*, *KIF14*, *NCAPG,* and *NUSAP1* were the most significantly upregulated genes in mPPGL (Suppl. Fig. 15A), thereby displaying an expression trend, which is opposite to what we observed in combination-treated PC12 cells *versus* control cells (Fig. 6E). This gene signature was also analyzed in the TCGA dataset, and only three among the mitosis-related genes were significantly upregulated in mPPGL *versus* non-metastatic tumors (i.e. *ASPM, CENPF, KIF14*), probably due to the low numbers of metastatic cases in this cohort (n= 16) (Suppl. Fig. 17A).

To gain further insight in the association between the mitotic gene signature and the metastatic phenotype, patients from the CNIO cohort were stratified into quartiles based on gene expression, and we looked for a correlation between expression levels and clinical behavior/characteristics of the tumors. We observed that samples with higher expression of these genes were overrepresented in aggressive/metastatic human PPGL when compared to non-metastatic cases (Fig. 6F). This phenomenon was especially pronounced for *ASPM*, *CENPF*, *CEP55*, *KIF14*, *NCAPG* and *NUSAP1* (Fig. 6F).

The mitotic gene signature was also checked in the RNAseq expression data from the CNIO cohort. An unsupervised clustering of PPGL samples revealed that not only samples from metastasis (black, tumor type), but also primary tumors that later developed metastasis (red, clinical outcome) are associated with a high expression of the 9-mitosis-signature gene list (Fig. 7A). Most of these cases belong to the C1-pseudohypoxia cluster, for which a higher risk for metastastic disease is well recognized [11,13]. Additionally, when we performed clustering of a series of primary tumors and their corresponding metastases derived from the same patients (paired samples) this 9-gene signature was more highly expressed in the metastases (Fig. 7B), albeit these genes were already upregulated in the primary PPGLs compared to the benign cases (Fig. 7A).

**Figure 7.**
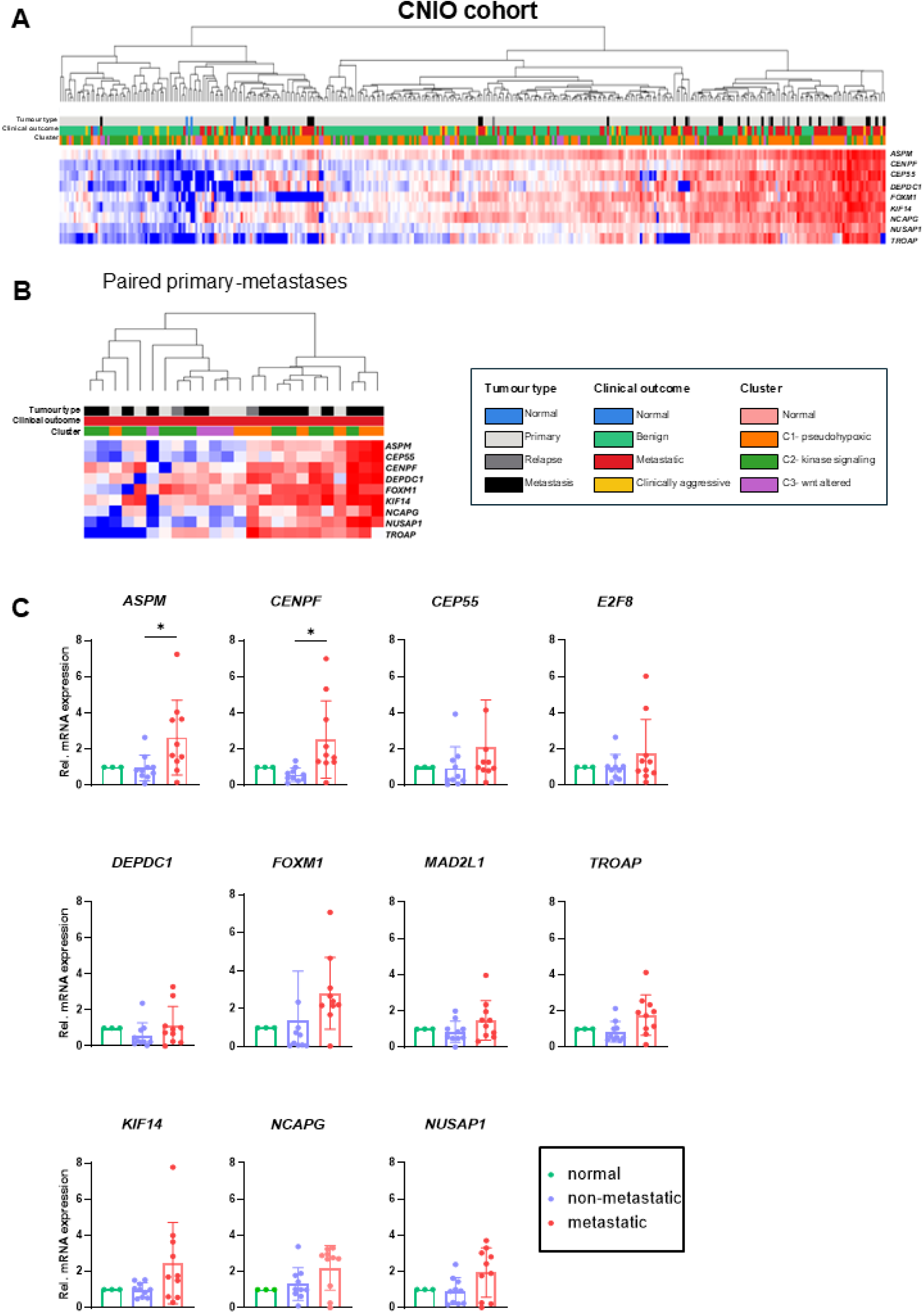
Mitotic 9-gene signature expression in mPPGLs. (**A-B**) Heatmap representation of gene expression in the CNIO cohort, displaying unsupervised hierarchical clustering of tumor samples based on selected gene expression profiles. Tumor type, clinical outcome, and molecular clustering information are annotated. Total cohort samples (**A**) and primary-metastatic pairs illustrating differences in expression between primary tumors and their metastatic counterparts (**B**) were analyzed. (**C**) Scatter plots depicting qRT-PCR results of the selected genes across non-metastatic and metastatic samples from primary human PPGL tissues. Each dot represents an individual sample, with error bars indicating variability. Statistics: 1way ANOVA. Only significant comparisons are shown. *, p< 0.05.

The expression of the mitosis-related genes was further validated by analyzing 10 metastatic and 10 non-metastatic independent PPGLs by realtime qRT-PCR. These analyses again confirmed the higher expression of *NUSAP1*, *CEP55*, *DEPDC1*, *TROAP, KIF14, NCAPG, ASPM, CENPF*, *MAD2L1* in mPPGLs compared with the non-metastatic cases (Fig. 7). Altogether, this suggests that mPPGLs upregulate proteins involved in mitotic spindle assembly to ensure DNA replication, likely representing an oncogenic dependency, while dual PI3K and CDK4/6 inhibition effectively downregulates these genes.

## DISCUSSION

Unresectable mPPGLs remain a significant therapeutic challenge due to their association with substantial morbidity and elevated mortality. By harnessing the molecular mechanisms driving PPGL progression, we assessed the therapeutic potential of targeting two essential processes, i.e. PI3K signaling and cell cycle regulation. Our study shows that combined pan-PI3K (buparlisib) and CDK4/6 (ribociclib) inhibition induces strong antitumor effects, particularly *in vivo*. This efficacy results from the downregulation of genes involved in mitotic spindle assembly and chromosomal segregation, a response unique to the dual treatment. Given that this gene signature is upregulated in metastatic versus non-metastatic human PPGLs, these tumors may be especially sensitive to dual PI3K + CDK4/6 targeting, thereby defining a potential therapeutic vulnerability.

Mechanistically, PI3K and CDK4/6 inhibition impinge on FOXM1 reducing its expression and activity, a feature unique of combination-treated PPGL cells. FOXM1, a master regulator of mitosis and oncogenic driver, controls genes critical for cell cycle progression, apoptosis, and DNA repair [57]. Upregulation of FOXM1 is common in cancer and helps tumor cells manage aneuploidy, a feature of aggressive cancers like advanced PPGLs [38]. Furthermore, FOXM1 overexpression leads to tolerance against chromosomal instability induced by replication errors, thereby inducing chemotherapy resistance [58–60]. In contrast, FOXM1 downregulation impairs mitosis, increases spindle abnormalities, and induces tumor cell death [52,53,61]. Thus, mPPGLs and, more broadly, tumors characterized by elevated FOXM1 activity, are expected to be particularly susceptible to therapeutic strategies, such as dual PI3K-CDK4/6 inhibition, that disrupt the FOXM1-mediated regulation of proper mitotic spindle formation.

FOXM1 is a key transcription factor implicated in multiple tumor types, making it an attractive therapeutic target and driving extensive efforts to develop effective inhibitors. However, current FOXM1 inhibitors face significant challenges, including limited potency, poor specificity, and potential off-target effects [62]. FOXM1 activation is regulated by multiple converging oncogenic pathways, such as Ras/Raf/ERK, Myc, HIF and PI3K/mTOR, known to play important roles in PPGL progression [63,64]. Our strategy of targeting both the PI3K and CDK4/6 pathways simultaneously effectively prevented compensatory FOXM1 activation by alternative upstream regulators. Notably, FOXM1 inhibitors are also used in combination therapies to enhance efficacy and mitigate resistance mechanisms commonly observed with monotherapies [65]. In breast cancer, a proposed therapeutic approach combines a FOXM1 inhibitor with a CDK4/6 inhibitor [66], suggesting potential synergistic effects that extend beyond FOXM1 inhibition alone.

Mining RNASeq data from the CNIO human PPGL cohort and from the TCGA-PPGL dataset revealed significant upregulation of the mitotic spindle category in metastatic versus non-metastatic tumors, more pronounced in the CNIO cohort having more metastatic cases [19,38]. GSEA analyses highlighted a positive enrichment of gene sets linked to cell cycle regulation, including E2F targets, G2M checkpoint, and mitotic spindle assembly, with chromosome segregation processes upregulated and oxidative phosphorylation pathways downregulated in metastatic tumors. The mitotic spindle was also among the upregulated pathways in SDHx-mutated mPPGLs by Zethoven et al. [39]. Focusing on this pathway, we identified a 9-gene classifier, highly expressed in metastatic/relapsing PPGLs, especially but not exclusively belonging to the most aggressive pseudohypoxia cluster.

The CNIO dataset is an incredible valuable resource, as it includes primary tumor samples with known metastatic outcomes. Our signature gene list successfully distinguished not only metastatic from non-metastatic tumors, but also primary tumors that later developed metastases from those that did not, demonstrating its potential as an additional predictor of tumor aggressiveness. The same 9-gene-list was also increased in the metastatic TCGA samples and additionally in unrelated PPGL samples by qPCR, thus validating its usefulness over several patient cohorts. Prediction of metastatic risk for PPGLs is crucial to improve patients’ survival. Recently, a machine learning-based tool considering clinical phenotypes [67], and a molecular classifier taking into account microsatellite instability, somatic ATRX/TERT alterations, *CDK1* overexpression and MAML3-fusions have shown promise for predicting mPPGL prognosis [38]. Our study identifies a 9-gene signature associated with metastatic risk: it is overexpressed in primary PPGLs which later developed metastases, and also in metastases *versus* primary matched samples. While the 9-gene signature may not be sufficient as a single factor to predict metastatic behaviour, the mitotic signature list could prove to be useful together with other parameters (PASS Score, genetic driver mutation) to stratify patients into risk groups.

We asked ourselves whether the PI3K and CDK inhibitors that we used could be substituted with different drugs of the same class. For both targets several drugs have been developed and used in various tumor settings. The pan-PI3K inhibitor buparlisib that we have used in our study has proven efficacy in several studies. When combined with paclitaxel, buparlisib demonstrated improved clinical efficacy with a manageable safety profile in platinum-pretreated recurrent or metastatic head and neck squamous cell carcinoma (HNSCC) [68].

To further assess its therapeutic potential in this population, a Phase III trial (NCT04338399) is currently ongoing. Isoform-specific PI3K inhibitors have also shown promise: alpelisib, targeting PI3Kα, is approved with fulvestrant for PIK3CA-mutant, ER-positive breast cancer [69], and duvelisib, a PI3Kδ inhibitor, is approved for relapsed/refractory chronic lymphocytic leukemia or small lymphocytic lymphoma [70,71]. We previously demonstrated that alpelisib, combined with the mTOR inhibitor everolimus, inhibits the viability of human primary PPGL cells [24], but we now observed no increased antitumor efficacy by combining it with CDK4/6 inhibition. In contrast, duvelisib was effective when administered together with ribociclib in PC12 cells. In our setting, only pan-PI3K or PI3Kδ inhibition had synergistic effects in the combination of CDK4/6 inhibition. It will be very interesting to identify the detailed mechanism of this interplay.

CDK4/6 inhibitors combined with aromatase inhibitors are approved as first-line treatment for HR+/HER2-advanced breast cancer, significantly improving progression-free and overall survival [72–74]. Ribociclib has also been approved as adjuvant therapy for early high-risk breast cancer [75]. The available CDK4/6 inhibitors, i.e. palbociclib, ribociclib, and abemaciclib differ in pharmacologic properties, toxicity, and resistance mechanisms [76], necessitating careful patient stratification for optimal outcomes. In line with this finding, in the rat PC12 cell line, ribociclib, but not palbociclib, nor abemaciclib, enhanced the anti-proliferative effects of buparlisib.

Unfortunately, there are no rat specific antibodies for all relevant FOXM1 phosphorylation sites available and no representative human PPGL cell lines to dive further into this topic. However, it will be helpful in future to expand the spectrum of effective drug combinations with the help of primary patient derived cultures to overcome therapy resistance and improve outcomes by patients’ stratification.

Previous preclinical studies have also shown the efficacy of dual CDK4/6 and PI3K inhibition against hormone receptor (HR)+ breast cancers and triple-negative breast cancer [77,78]. Additionally, the resistance of breast cancer cells to palbociclib can be overcome by the administration of PI3K inhibitors [79], further supporting a potential benefit of a combination therapy. Yet, our data provide additional new mechanistic insights to the mechanism of dual CDK4/6 and PI3K inhibition, highlighting unique molecular effects such as the downregulation of FOXM1 and mitotic spindle genes, and the disruption of correct chromosomal segregation. These findings suggest a broader impact of the combination therapy on cell cycle progression and genomic stability than previously recognized.

The superior efficacy of the combinatorial PI3Ki+CDK4/6i treatment compared to single-agent therapies was demonstrated, among others, in primary cultures derived from human tumors with diverse genetic backgrounds and anatomical origins, including metastatic cases, attesting to its broad applicability. This dual inhibition approach was further validated in PC12-derived xenografts, where it suppressed tumor growth and induced necrosis more effectively than single treatments. Notably, the combination remained highly effective even with reduced doses of buparlisib and ribociclib, suggesting potential for minimizing side effects in future clinical applications. The superior performance of the combination therapy aligns with the existing literature on the roles of PI3K and CDK4/6 pathways in cancer [80,81] and corroborates the centrality of these pathways in adrenomedullary tumorigenesis [16–18,38,39].

Altogether, the combined targeting of the PI3K and CDK4/6 pathways offers a compelling therapeutic strategy for treating metastatic PPGLs. By disrupting key mitotic processes and leveraging the vulnerabilities of rapidly dividing cancer cells, this approach demonstrates robust efficacy while potentially minimizing toxicity. The enhanced efficacy of the combination therapy can be attributed to its ability to induce mitotic spindle defects and trigger mitotic catastrophe. By causing irreversible damage, this approach not only impairs tumor growth but also minimizes the likelihood of tumor relapse, as it prevents the survival of cells capable of developing resistance. The findings presented here lay the groundwork for future studies focused on optimizing dosing regimens, reducing side effects, and translating this promising therapeutic strategy into clinical practice to address the unmet needs of patients with aggressive PPGLs currently having limited treatment options.

