## Supplemental Tables and Figures for "A genetic signature predicts aggressive paraganglioma sensitivity to dual PI3K-CDK4/6 inhibition therapy"

Karna et al.

Supplementary Tables (n=5)

**Supplementary Table 1: Drug concentrations used to determine the $IC_{50}$ of each compound**

| PC12 cells | | |
|---|---|---|
| **CDK4/6i [μM]** | **PI3Ki [μM]** | **Combo [μM]** |
| 0.1 | 0.1 | 0.01 + 0.04 |
| 0.2 | 0.2 | 0.05 + 0.2 |
| 0.5 | 0.5 | 0.1 + 0.4 |
| 1 | 1 | 0.2 + 0.8 |
| 5 | 2 | 0.5 + 2 |
| 10 | 4 | 1 + 4 |
| 20 | 8 | 2 + 8 |

| MPC cells | | |
|---|---|---|
| **CDK4/6i [μM]** | **PI3Ki [μM]** | **Combo [μM]** |
| 0.2 | 0.05 | 0.05 + 0.2 |
| 0.4 | 0.1 | 0.1 + 0.4 |
| 0.8 | 0.2 | 0.2 + 0.8 |
| 2 | 0.5 | 0.5 + 0.2 |
| 4 | 1 | 1 + 4 |
| 8 | 2 | 2 + 8 |
| 16 | 4 | 4 + 16 |

**Supplementary Table 2: Summary of human patient PPGLs used in this study.**

This table summarizes the patient demographic and tumor characteristics of samples used in the study (Figure 2 G,H). PCC, pheochromocytoma; PGL, paraganglioma.

| Sample Name | Patient Age (years) | Patient Sex | Tumor Type/ Histology | Tumor size (cm) | PPGL Cluster | Anatomical Site | Sample Type | Passage Number | Biochemical phenotype | Key Genetic Alterations | |
|---|---|---|---|---|---|---|---|---|---|---|---|
| | | | | | | | | | | Somatic | Germline |
| Patient 1 | 70 | Female | PGL | 4,2 | Pseudohypoxia | Abdominal | Primary Tumor | P0 | Silent | Negative | Negative |
| Patient 2 | 19 | Female | PCC | 6,1 | Pseudohypoxia | Adrenal | Primary Tumor | P0 | Noradrenergic | *VHL* | Negative |
| Patient 3 | 34 | Female | PGL | 4,5 | Pseudohypoxia | Abdominal | Primary Tumor | P0 | Noradrenergic | Negative | *VHL* |
| Patient 4 | 53 | Male | PCC, Metastatic | 5,2 | Undefined, aggressive | Adrenal | Primary Tumor | P15 | Noradrenergic | *ATRX* | Negative |
| Patient 5 | 62 | Male | PCC | 6,7 | Kinase signaling | Adrenal | Primary Tumor | P0 | Noradrenergic | *TMEM127* | Negative |
| Patient 6 | 65 | Female | Metastasized PCC | n/a | Kinase signaling, aggressive | Spine | Metastatic site | P0, P5 | Noradrenergic | *ATRX*, 20% *cMET* vus | *NF1* |

| Supplementary Table 3: Taqman Assays used in this study (Ref: Thermo Fischer Scientific inventory) | | | |
|---|---|---|---|
| **Taqman assays** | | | |
| **Gene Name** | **Assay ID** | **Probe Type** | **Amplicon Length** |
| *Aspm* | Rn01758905_m1 | FAM-MGB | 60 |
| *ASPM* | Hs00411505_m1 | FAM-MGB | 77 |
| *Ccna1* | Rn01761348_m1 | FAM-MGB | 102 |
| *Ccna1* | Mm00432337_m1 | FAM-MGB | 59 |
| *Ccna2* | Rn01493715_m1 | FAM-MGB | 107 |
| *Ccna2* | Mm00438063_m1 | FAM-MGB | 83 |
| *Cenpf* | Rn01411515_m1 | FAM-MGB | 75 |
| *CENPF* | Hs01118845_m1 | FAM-MGB | 77 |
| *Cep55* | Rn01479711_m1 | FAM-MGB | 122 |
| *CEP55* | Hs01070181_m1 | FAM-MGB | 76 |
| *Depdc1* | Rn01521456_m1 | FAM-MGB | 73 |
| *DEPDC1* | Hs00873600_g1 | FAM-MGB | 71 |
| *E2f8* | Rn01476915_m1 | FAM-MGB | 57 |
| *E2F8* | Hs00226635_m1 | FAM-MGB | 68 |
| *Foxm1* | Rn00668556_m1 | FAM-MGB | 86 |
| *FOXM1* | Hs01073586_m1 | FAM-MGB | 77 |
| *Kif14* | Rn01490403_m1 | FAM-MGB | 68 |
| *KIF14* | Hs00978236_m1 | FAM-MGB | 66 |
| *Mad2l1* | Rn01425146_m1 | FAM-MGB | 89 |
| *MAD2L1* | Hs01554513_g1 | FAM-MGB | 65 |
| *Ncapg* | Rn01443626_m1 | FAM-MGB | 72 |
| *NCAPG* | Hs00254617_m1 | FAM-MGB | 109 |
| *Nusap1* | Rn01515209_g1 | FAM-MGB | 74 |
| *NUSAP1* | Hs01006195_m1 | FAM-MGB | 87 |
| *RPS18* | Hs01375212_g1 | VIC-MGB_PL | 93 |
| *Rps18* | Rn01428913_gH | VIC-MGB_PL | 62 |
| *Rps18* | Mm02601777_g1 | VIC-MGB_PL | 76 |
| *Troap* | Rn01532712_m1 | FAM-MGB | 79 |
| *TROAP* | Hs04400287_g1 | FAM-MGB | 62 |

**Supplementary Table 4: Antibodies used in this study and their dilutions**

**Antibodies**

| Type | Antibody | Clonality | Company | ID | Dilution |
|---|---|---|---|---|---|
| Primary | Phospho-S6 Ribosomal Protein (Ser240/244) (D68F8) XP® | Rabbit | Cell Signaling Technology | 5364 | 1/2500 |
| Primary | S6 Ribosomal Protein (5G10) | Rabbit | Cell Signaling Technology | 2217 | 1/2500 |
| Primary | Akt Antibody | Rabbit | Cell Signaling Technology | 9272 | 1/1000 |
| Primary | Phospho-Akt (Ser473) (D9E) XP® | Rabbit | Cell Signaling Technology | 4060 | 1/2000 |
| Primary | MAD2L1 Polyclonal antibody | Rabbit / IgG | Proteintech | 10337-1-AP | 1/1000 |
| Primary | α-Tubulin antibody, clone AA13 | Mouse | Sigma-Aldrich | T8203 | 1/3000 |
| Secondary | Rabbit IgG (H/L):HRP | Goat | Bio-Rad Laboratories | STAR208P | 1/2000 |
| Secondary | Mouse IgG (H/L):HRP | Goat | Bio-Rad Laboratories | STAR207P | 1/2000 |
| Secondary | Mouse IgG (H+L), F(ab')2 Fragment (Alexa Fluor® 647 Conjugate) | Goat | Cell Signaling Technology | 4410 | 1/2000 |

**Supplementary Table 5: Reagents, Equipments and Softwares used in this study**

**Reagents, Kits and Chemicals**

| Type | Company | Catalogue Number |
|---|---|---|
| RPMI1640 Medium | Thermo Fisher Scientific | 61870044 |
| BCA Protein Assay Kit | Thermo Fisher Scientific | 23225 |
| Buparlisib (PI3Ki) | MedChemExpress | HY-70063 |
| Caspase-Glo® 9 Assay | Promega | G8211 |
| CyQUANT® NF Cell Proliferation Assay | Thermo Fisher Scientific | C35006 |
| DMSO | Sigma-Aldrich | D2650 |
| Fast Advanced Master Mix | Thermo Fisher Scientific | 4444557 |
| Fetal Bovine Serum (FBS) | Thermo Fisher Scientific | A5256801 |
| High-Capacity RNA-to-cDNA Kit | Thermo Fisher Scientific | 4387406 |
| Horse Serum (HS) | Thermo Fisher Scientific | 26050088 |
| Matrigel | Corning | 356231 |
| Penicillin-Streptomycin | Thermo Fisher Scientific | 15140122 |
| RealTime-Glo™ Annexin V Apoptosis and Necrosis Assay | Promega | JA1011 |
| RealTime-Glo™ MT Cell Viability Assay | Promega | G9712 |
| Ribociclib (CDK4/6i) | MedChemExpress | HY-15777 |
| RNeasy Mini Kit | Qiagen | 74104 |
| SuperSignal West Pico Chemiluminescent Substrate | Thermo Fisher Scientific | 34080 |
| Tumor Dissociation Kit (Human) | Miltenyi Biotec | 130-095-929 |
| Tumor Dissociation Kit (Mouse) | Miltenyi Biotec | 130-096-730 |

**Equipments**

| Equipment | Manufacturer | Model/Catalogue Number |
|---|---|---|
| Automated Immunostainer | Ventana Medical Systems | BenchMark XT |
| Confocal Microscope | Olympus | FV1200 |
| gentleMACS™ Octo Dissociator | Miltenyi Biotec | 130-096-427 |
| Inverted Routine microscope | Nikon | Eclipse *TS* 2 |
| Microplate reader | Thermo Fischer Scientific | Varioscan LUX |
| NanoDrop | Thermo Fischer Scientific | ND-1000 |
| Pathology scanner | Leica Biosystems | Aperio AT2 |
| Ultra-Low Attachment Plates (96-well) | Corning | 3474 |
| Upright Clinical Microscope | Nikon | Eclipse C*i* -L plus |
| X-ray Irradiator | Precision X-ray | X-RAD 320 |

**Software and Analysis Tools**

| Software | Version |
|---|---|
| ChEA3 | v3 |
| decoupleR | 2.9.7 |
| Enrichr | Online Tool |
| GraphPad Prism | 10 |
| GSEA | 4.3.3 |
| Image Lab, Bio-Rad | 6.1.0 |
| ImageJ | 1.53t, Java8 |
| Nextflow nf-core/rnaseq | 3.14.0 |
| QuPath | 0.4.4 |
| R-Project | R-4.4.2 |
| STRING Database | 12 |

Karna et al.

Supplementary Figures (n=18)

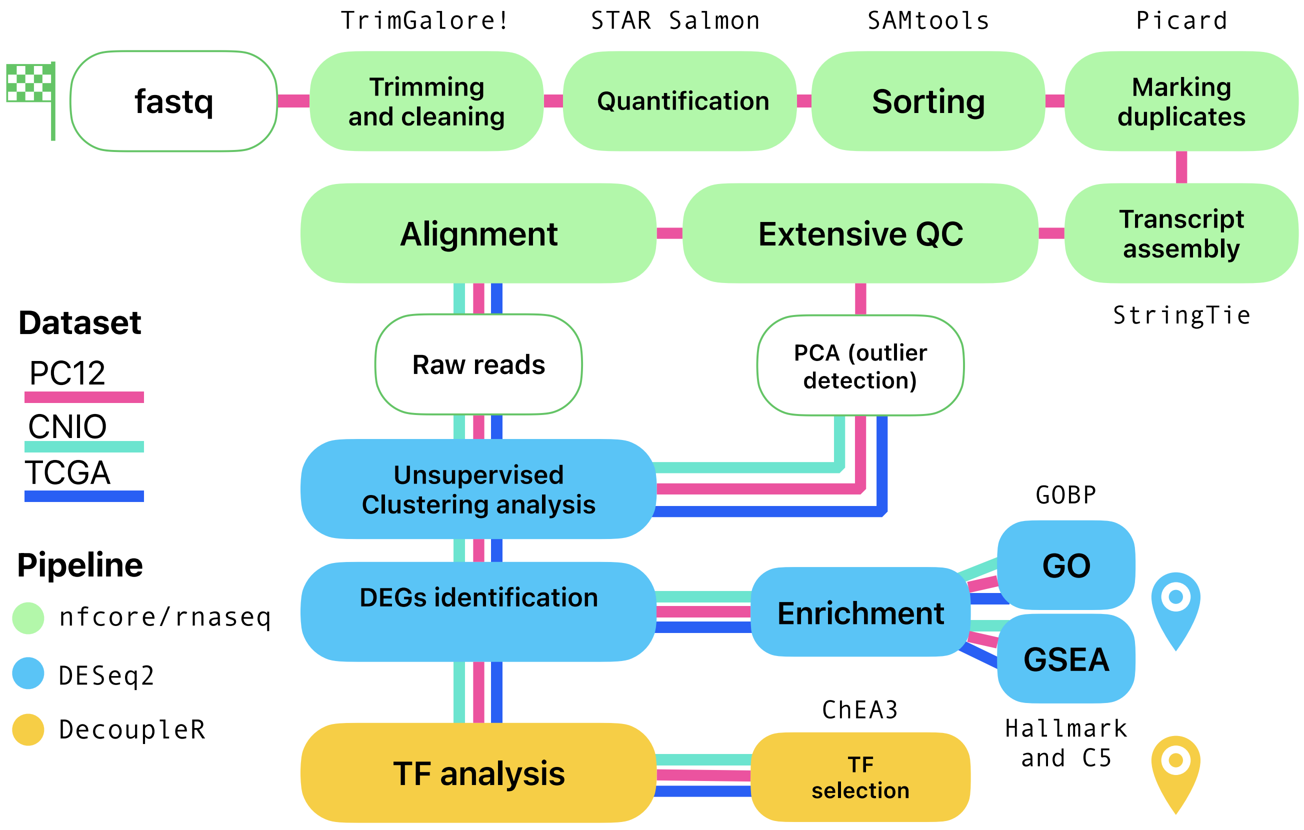


**Supplementary Figure S1.** Scheme of the workflow used for the processing of the RNAseq raw counts of the PC12 cells treated with CDK4/6i, PI3Ki or their combination (as indicated in the M&Ms). The workflow followed three steps: in the first one (green) the Nextflow nf-core/rnaseq pipeline was used to perform trimming, multiple alignment, sorting, marking duplicates, assembly, extensive data quality control and mapping of the fastq files, to obtain the raw counts. In the second step (blue) the DESeq2 pipeline was used in R to process the reads and obtain the differentially expressed genes (DEGs); then the enrichment analysis was performed using both Gene Ontology (GO) and Gene Set Enrichment Analysis (GSEA). In the third step (yellow) the DecoupleR algorithm and the ChEA3 tool were used to analyze and to select the transcription factors (TFs). The Nextflow nf-core/rnaseq pipeline was used only to process fastq files from PC12 samples. The DESeq2 and the TF analysis were used on PC12 samples, CNIO samples and TCGA samples. Next to the different steps, the main used tools are indicated. The lines indicate the path followed by each cohort. DEG, differentially expressed genes; GSEA, gene set enrichment analysis; GO, gene ontology; QC, quality control; TF, transcription factor.

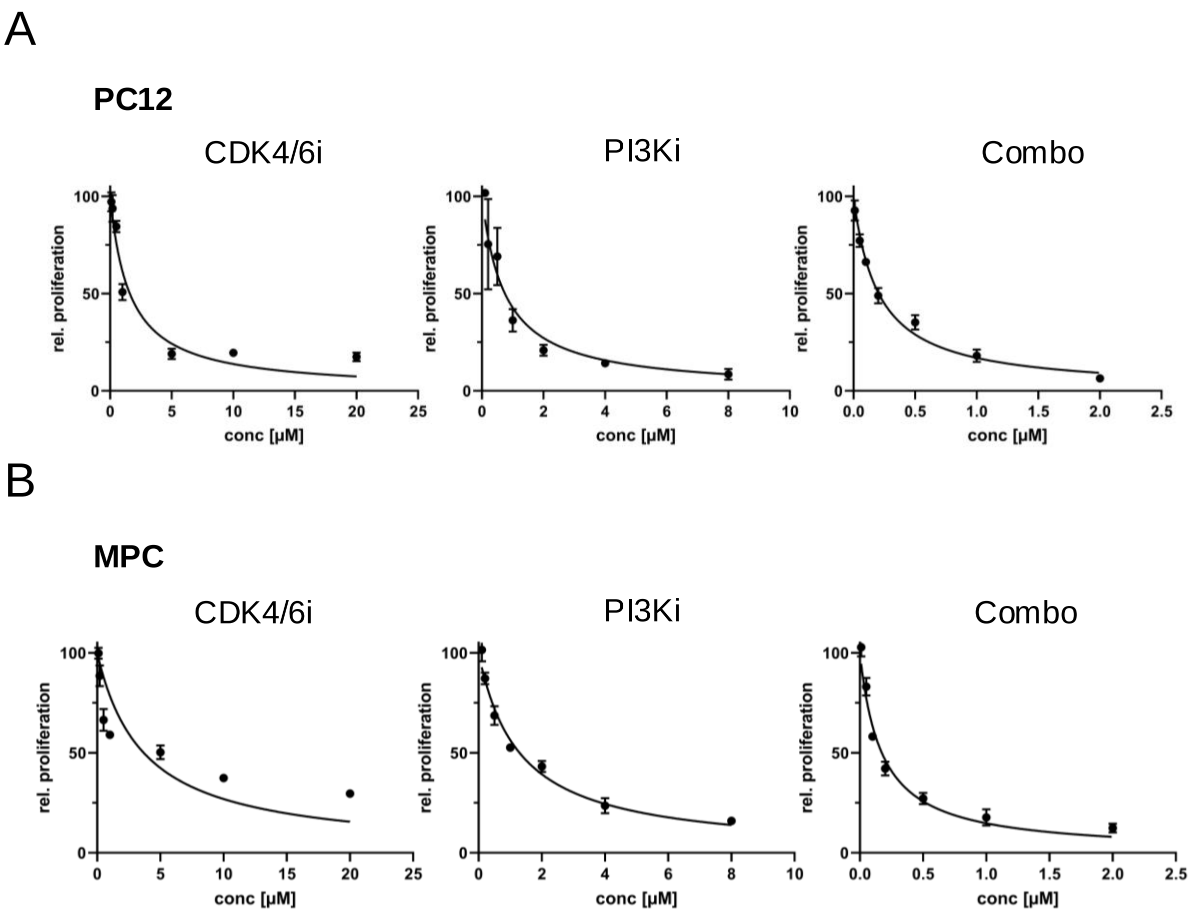


**Supplementary Figure S2.** Effect of CDK4/6i and PI3Ki treatment on the proliferation of PPGL cells. **(A,B)** Individual $IC_{50}$ curves of PC12 (**A**) or MPC (**B**) cells grown as 2D cultures and treated with CDK4/6i, PI3Ki, their combination or DMSO (control) (the combined curve for the treatments together are shown in Figure 1). Cell proliferation was measured 72h after treatment. The drug concentrations used to determine the $IC_{50}$ values are reported in Supplementary Table S1. The DMSO control was set to 100% and nonlinear regression was used to calculate the $IC_{50}$. Data shown are the mean ±SD from 3 independent experiments with 3 technical replicates each.
.

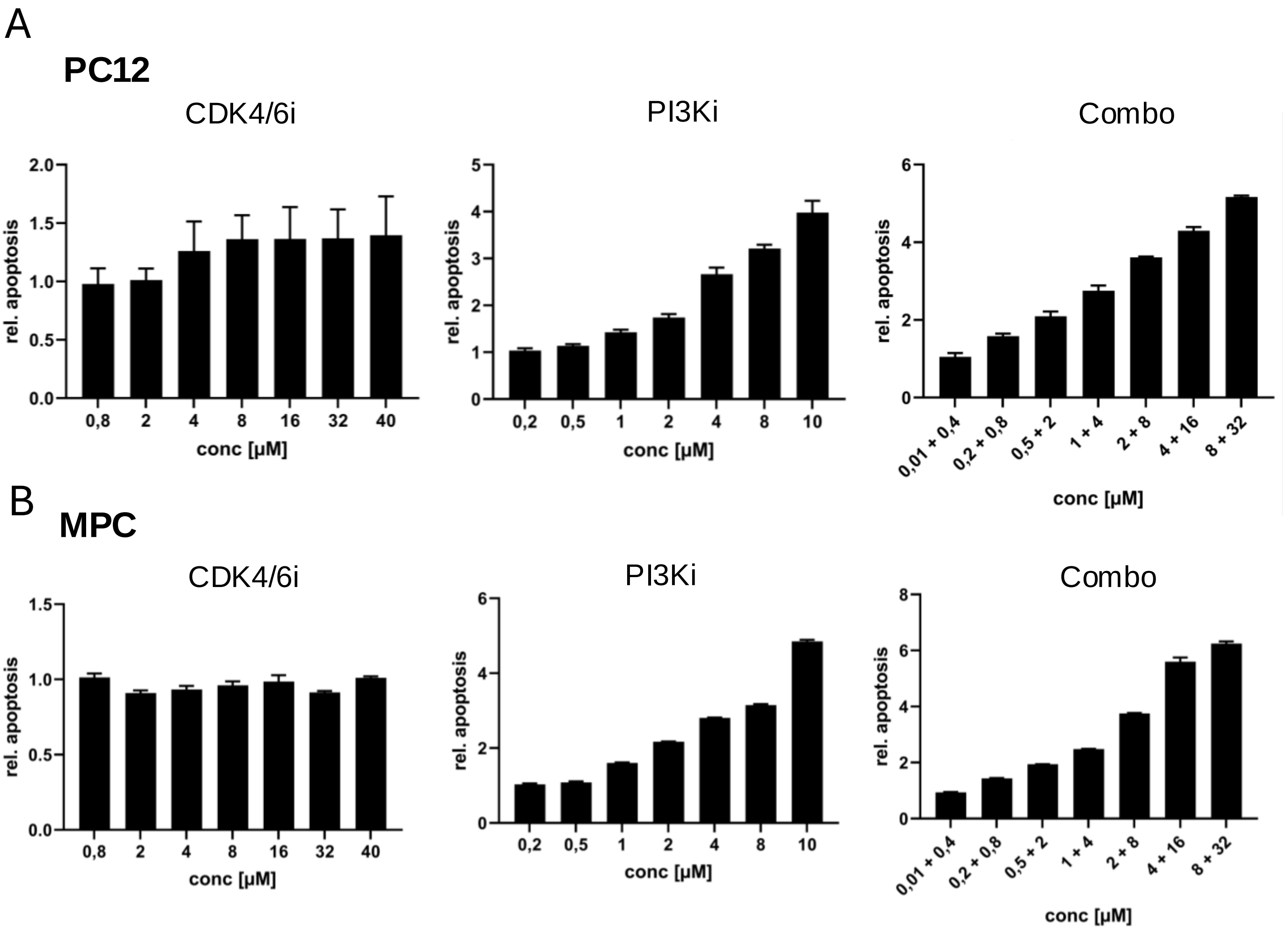


**Supplementary Figure S3**: Effect of PI3Ki and CDK4/6i on the induction of apoptosis of PPGL cells. **(A,B)** Induction of apoptosis for the individual drugs tested on PC12 **(A)** or MPC **(B)** cells (the treatments together are shown in Figure 1). Caspase 9 activity was assessed 72h after treatment with the indicated concentrations of the drugs alone or in combination, or with DMSO (control). Shown is the relative apoptosis normalized to the DMSO control. Data shown is the mean ± SD from 3 independent experiments with 3 technical replicates each. Statistics: 2way ANOVA.

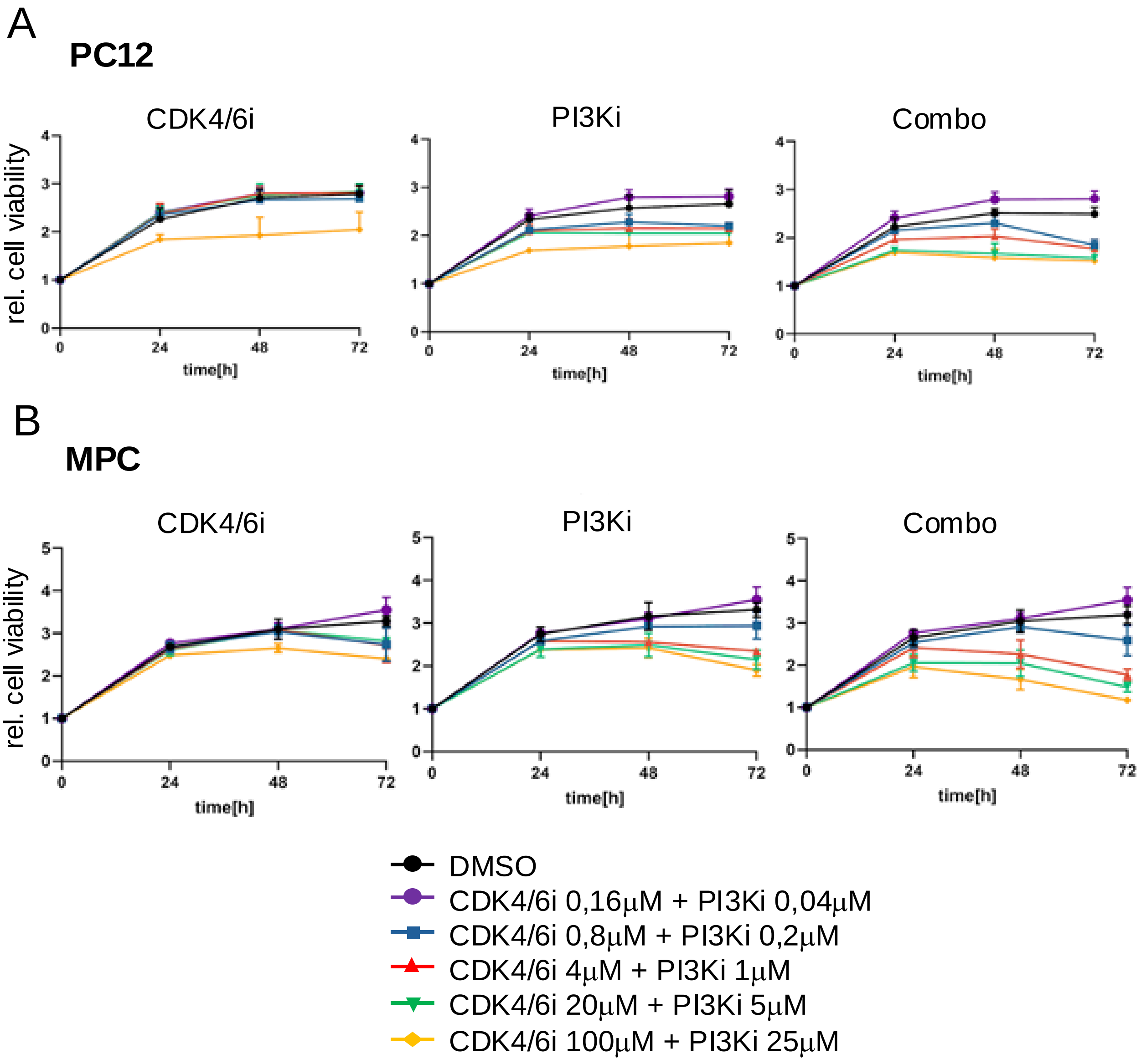


**Supplementary Figure S4**: Effect of PI3Ki and CDK4/6i on the viability of 3D organotypic cultures of PPGL cells. **(A,B)** Following spheroid formation, PC12 **(A)** or MPC **(B)** cells were treated with the indicated doses of CDK4/6i, PI3Ki, their combination or DMSO (control). Cell viability was assessed at time 0, 24h, 48h and 72h after treatment. Reported is the relative cell viability normalized against DMSO control (arbitrarily set to 1) and time 0. Data shown are the mean ±SD from 3 independent experiments with 8 technical replicates each. Statistics: 1way ANOVA.

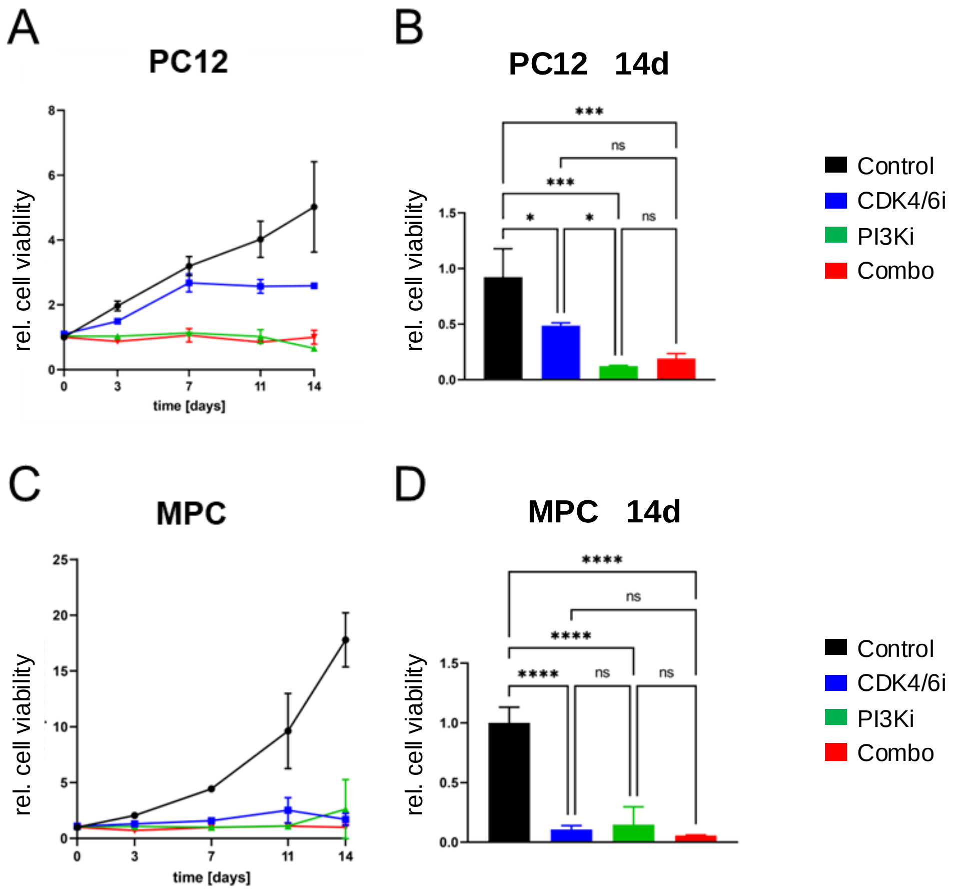


**Supplementary Figure S5**: Effect of PI3Ki and CDK4/6i on the spheroid size of PPGL cells grown as 3D organotypic cultures. **(A)** PC12 cells were plated in Gravity Plates and, after spheroid formation, treated with DMSO (control) or with the indicated drugs at the concentrations: CDK4/6i: PI3Ki: alone or in combination (Combo) for a total of 14 days. Spheroid size was measured at day 0, 3, 7, 11 and 14, and normalized against the size at day 0 (arbitrarily set to 1). **(B)** Spheroid size of PC12 cells treated as in A at day 14. **(C)** MPC cells were plated in Gravity Plates and, after spheroid formation, treated as in A for a total of 14 days. Spheroid size was measured at day 0, 3, 7, 11 and 14, and normalized against the size at day 0 (arbitrarily set to 1). **(D)** Spheroid size of MPC cells treated as in C at day 14. Data shown are the mean ±SD from 3 independent experiments with 8 technical replicates each. Statistics: 1way ANOVA.

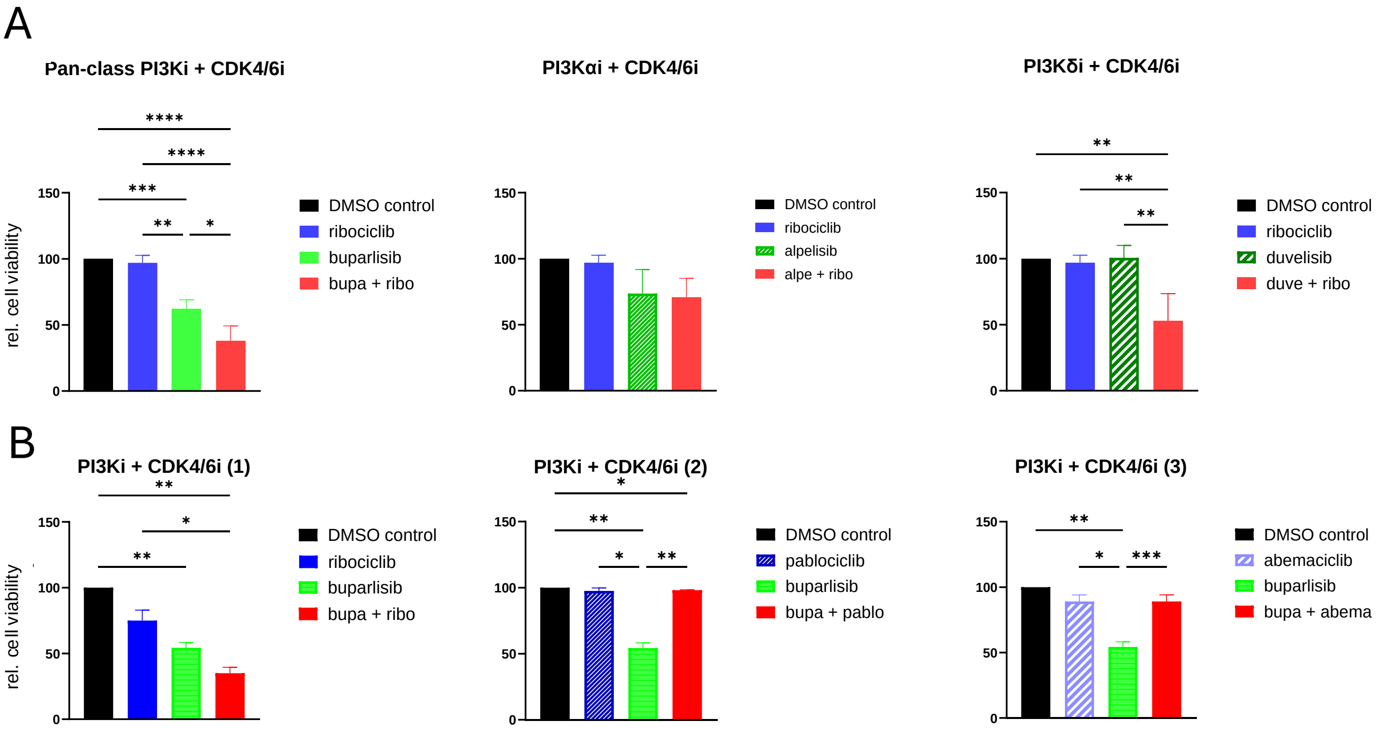


**Supplementary Figure S6**: Effect of alternative PI3K and CDK4/6 inhibitors on the proliferation of PPGL cells. **(A)** PC12 cells grown as 2D cultures were treated with IC50 of buparlisib, alpelisib and duvelisib alone or in combination with ribociclib, or with DMSO (control). Cell proliferation was measured 72h after treatment. The DMSO control was set to 100%. **(B)** PC12 cells grown as in A were treated with IC50 of ribociclib, palbociclib and abemaciclib alone or in combination with ribociclib, or with DMSO (control). Data shown are the mean ±SD from 3 independent experiments with 3 technical replicates each. ns, not significant; *, $p < 0.05$; **, $p < 0.01$; ***, $p < 0.001$; ****, $p < 0.0001$.

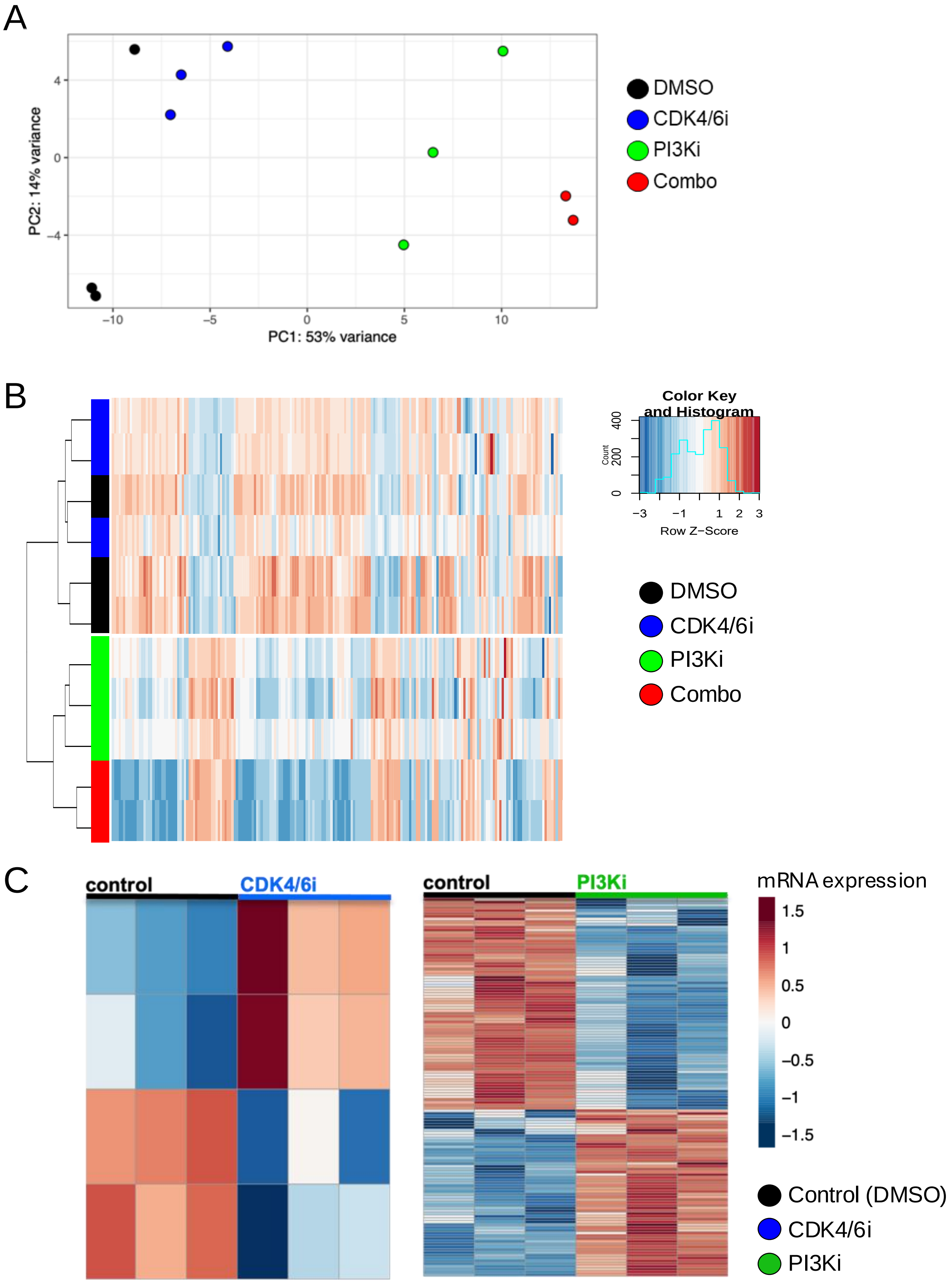


**Supplementary Figure S7**: **A)** PCA plot showing distribution of samples based on variation in gene expression. **B)** Unsupervised hierarchical clustering heatmap showing the expression of the top 200 most variable genes. DMSO = control, PI3Ki = buparlisb, CDK4/6i = ribociclib, Combo = buparlisib + ribociclib. All graphs were generated using R software (v 4.4.0). **C)** Heatmaps displaying the distribution of DEGs for control *vs* CDK4/6i and control *vs* PI3Ki comparisons. Red is for high mRNA expression levels and blue is for low mRNA expression levels. All graphs were generated using R software (v 4.4.0).

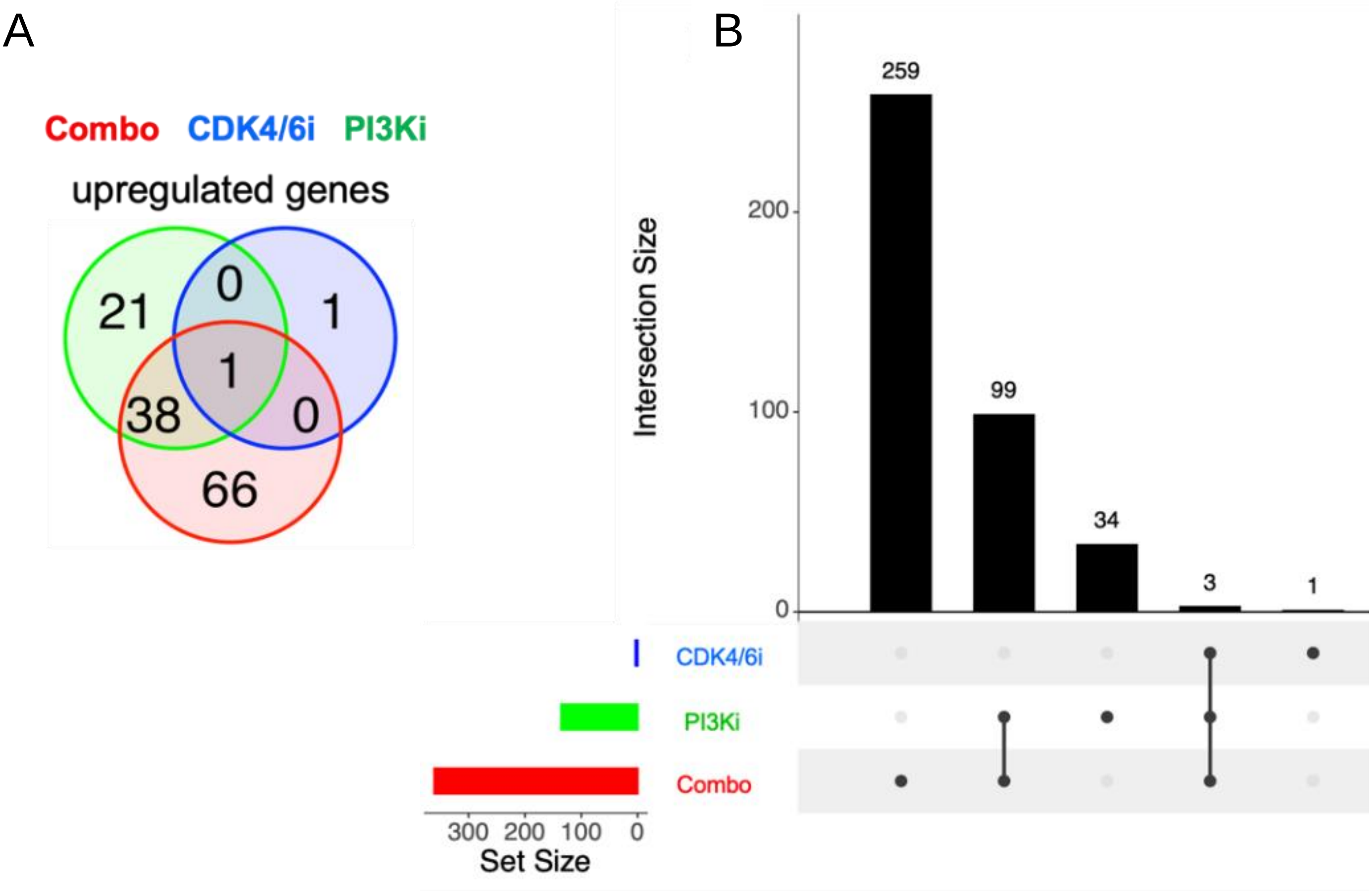


**Supplementary Figure S8**: **A**) Venn diagrams showin8 the intersection of up-regulated DEGs among the different treatment groups. **B**) Upset plot showing the overall DEG intersection among treatments groups. Set size = total number of DEGs for each condition; Intersection size = number of genes for each intersection. All graphs were generated using R software (v 4.4.0).

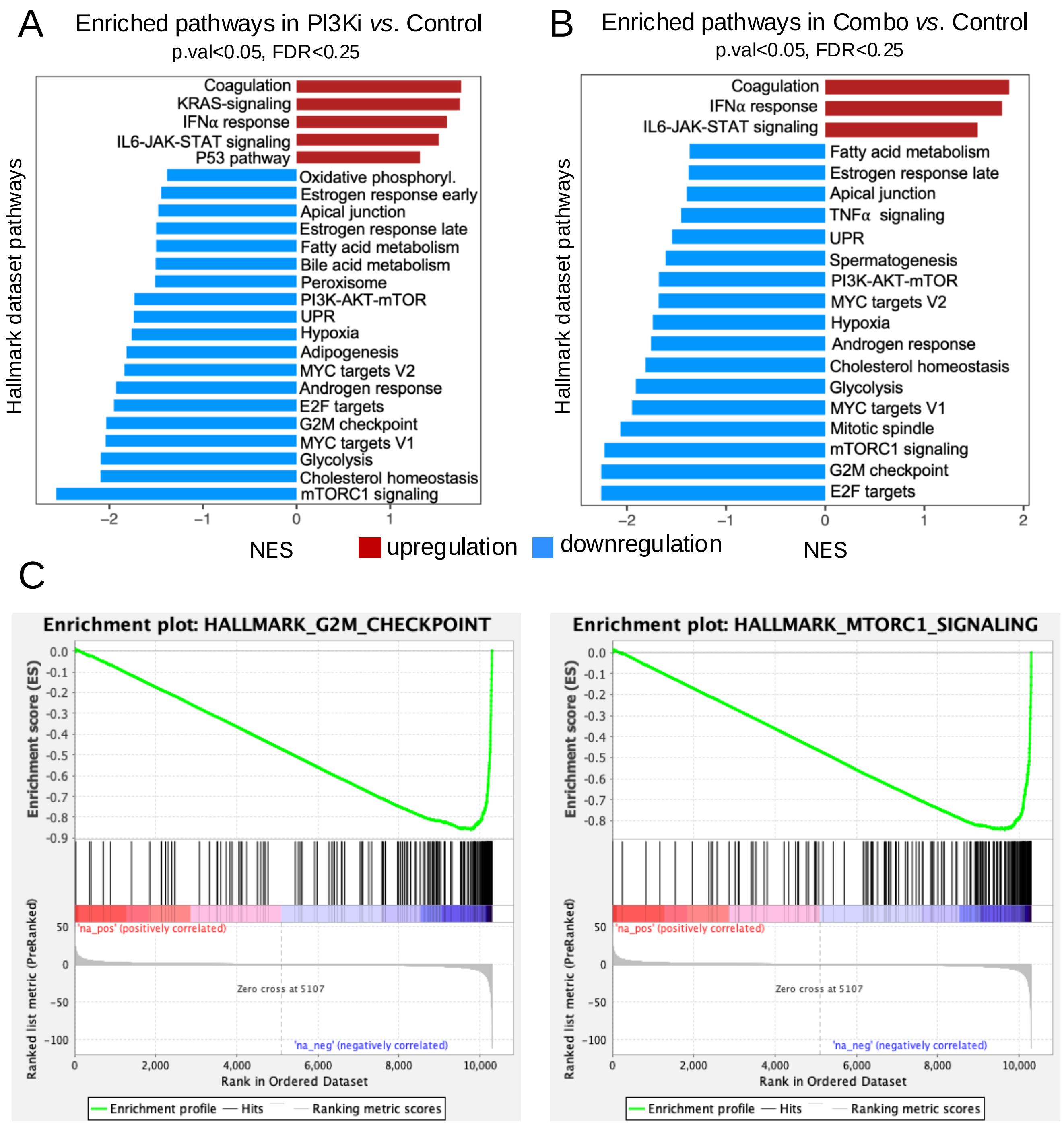


**Supplementary Figure S9. (A, B)** Bar plots showing the normalized enriched score (NES) relative to the PI3Ki *vs.* control **(A)** and combination *vs.* control **(B)** GSEA. **C)** GSEA plots showing the downregulated enrichment for the G2M checkpoint and MTORC1 signaling pathways. The horizontal axis represents the genes ranked by their expression levels, from most upregulated to most downregulated in a specific experimental condition. The vertical axis shows the cumulative Enrichment Score (ES), indicating how much a particular gene set is enriched (i.e., overrepresented) at the top, middle, or bottom of the ranked gene list. The main curve displays the ES trend across the X-axis. The downward peak indicates enrichment at the bottom of the ranked list (typically downregulated genes). The vertical black ticks along the X-axis indicate the positions of individual genes from the set within the ranked list. The density and distribution of these lines help visualize where the genes from the set are concentrated within the ranked dataset. The plots are generated using R software (v4.4.0). IL6 = interleukin 6; IFN $\alpha$ = interferon $\alpha$; IFN $\gamma$ = interferon $\gamma$; TNFA = tumor necrosis factor A; EMT = epithelial-mesenchymal-transition; UPR= unfolded protein response.

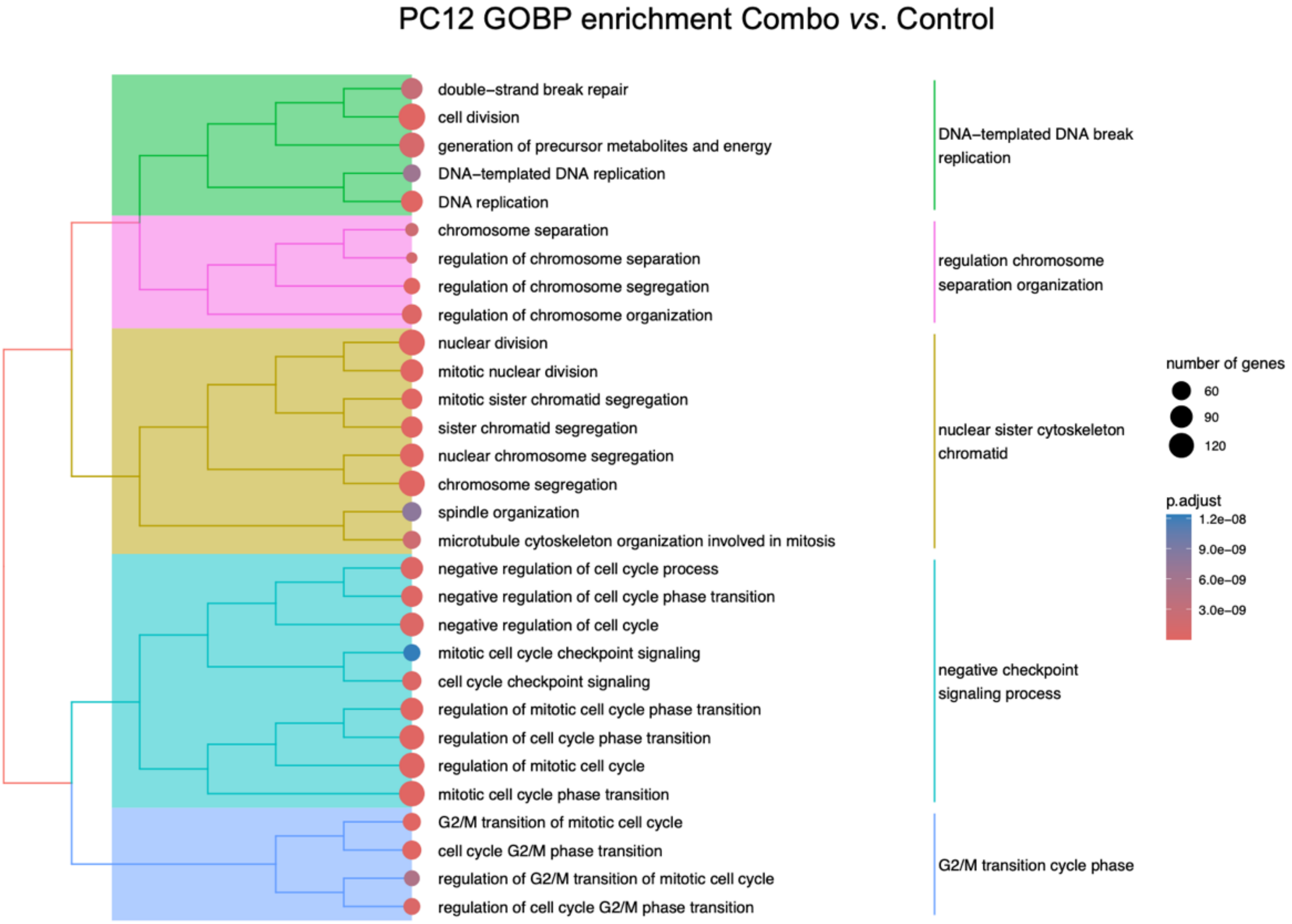


**Supplementary Figure S10**: Tree plot illustrating the enrichment of combination *vs.* control samples on GOBP category, highlighting significant processes that are grouped based on their functional similarities. Each node in the plot represents an enriched GO term, with the size of the node corresponding to number of genes associated with that term. Nodes that are closer together indicate related biological processes, while larger clusters represent broader functional categories. The tree branches representing related GO terms are shown in colors for easy identification. The color gradient of the nodes reflects the adjusted p-value, with red indicating higher statistical significance, as indicated on the right. The graph was generated using R software (v 4.4.0).

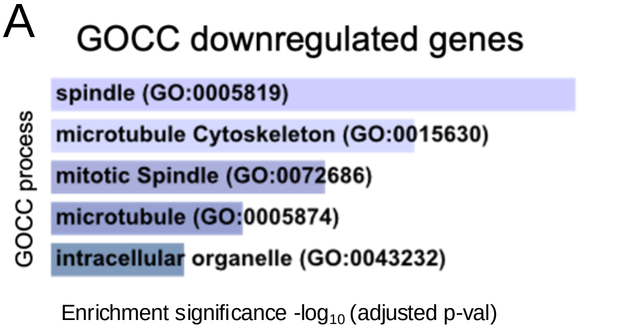


| GOCC term code | p-value | adj. p-value |
|---|---|---|
| GO:0005819 | 1.276e-20 | 1.416e-18 |
| GO:0015630 | 2.404e-15 | 1.334e-13 |
| GO:0072686 | 2.009e-12 | 7.432e-11 |
| GO:0005874 | 9.979e-10 | 2.769e-8 |
| GO:0043232 | 8.255e-8 | 0.000001833 |

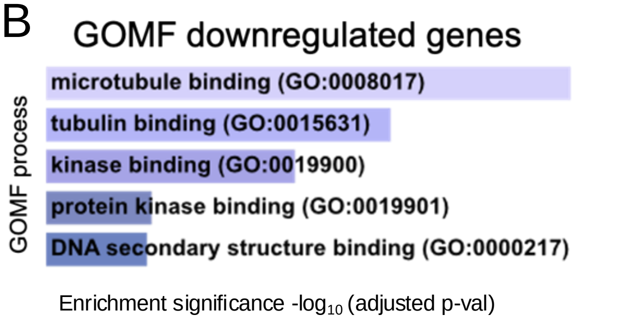


| GOMF term code | p-value | adj. p-value |
|---|---|---|
| GO:0008017 | 6.765e-11 | 1.786e-8 |
| GO:0015631 | 4.925e-8 | 0.000006501 |
| GO:0019900 | 0.00000163 | 0.0001440 |
| GO:0019901 | 0.0003106 | 0.01938 |
| GO:0000217 | 0.0003671 | 0.01938 |

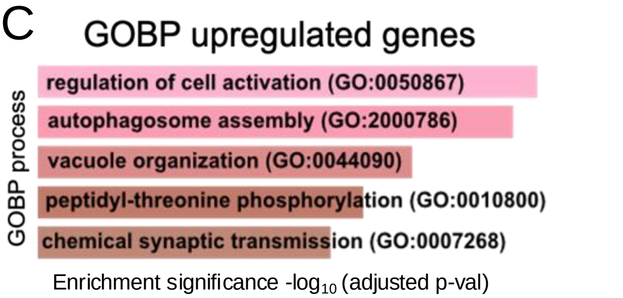


| GOBP term code | p-value | adj. p-value |
|---|---|---|
| GO:0050867 | 0.002024 | 0.3178 |
| GO:2000786 | 0.002306 | 0.3178 |
| GO:0044090 | 0.003978 | 0.3178 |
| GO:0010800 | 0.005182 | 0.3178 |
| GO:0007268 | 0.007825 | 0.3178 |

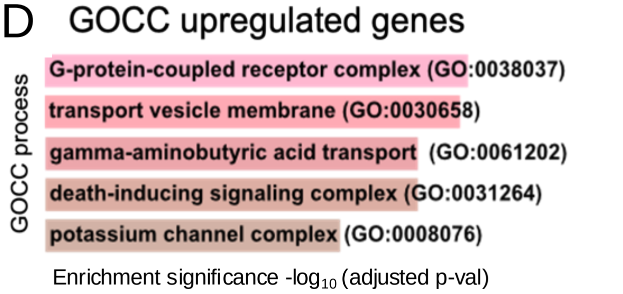


| GOBP term code | p-value | adj. p-value |
|---|---|---|
| GO:0038037 | 0.02670 | 0.3730 |
| GO:0030658 | 0.02994 | 0.3730 |
| GO:0061202 | 0.03108 | 0.3730 |
| GO:0031264 | 0.03108 | 0.3730 |
| GO:0008076 | 0.04282 | 0.4216 |

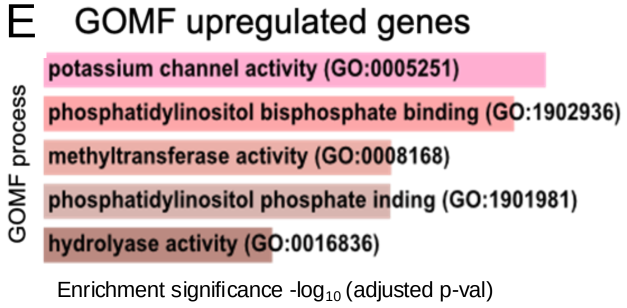


| GOMF term code | p-value | adj. p-value |
|---|---|---|
| GO:0005251 | 0.007015 | 0.2393 |
| GO:1902936 | 0.008542 | 0.2393 |
| GO:0008168 | 0.01204 | 0.2393 |
| GO:1901981 | 0.01313 | 0.2393 |
| GO:0016836 | 0.01823 | 0.2393 |

**Supplementary Figure S11**: **(A-E)** Bar plots indicating the top five enriched processes of genes wich are downregulated (A-B) and upregulated (C-E) uniquely in the combination *versus* control treatment, obtained using GOCC (A,D) and GOMF (B,E), and GOBP (C). On the right, the tables indicate the GO term code associated to p- and adjusted p-values. The analysis was conducted using Enrichr.

A

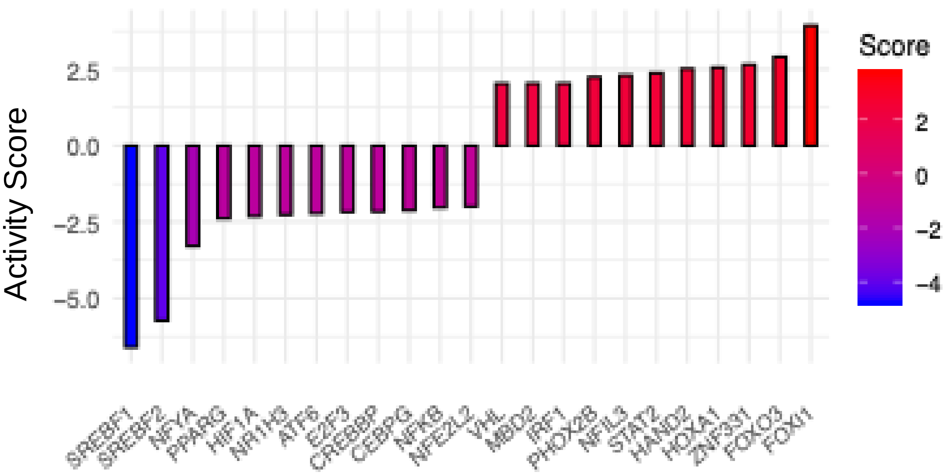


B

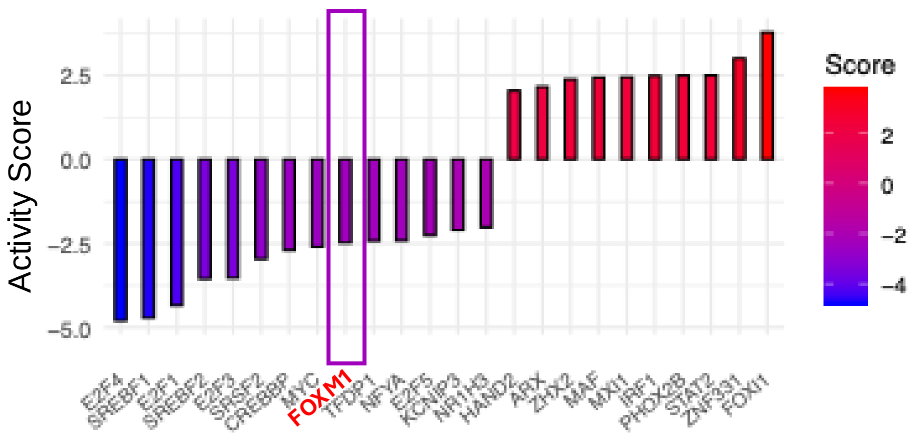


C ChEA 2022 transcription analysis

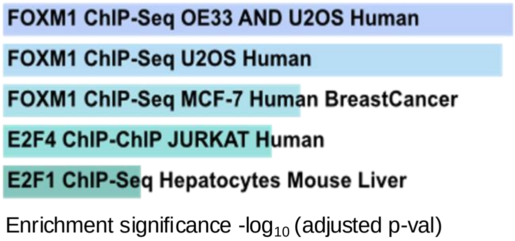


| GOBP term code | p-value | adj. p-value |
|---|---|---|
| FOXM1 25889361 | 5.129e-52 | 3.739e-49 |
| FOXM1 23109430 | 5.078e-51 | 1.851e-48 |
| FOXM1 26456572 | 2.405e-32 | 5.845e-30 |
| E2F4 17652178 | 1.053e-29 | 1.920e-27 |
| E2F1 26619117 | 1.387e-17 | 2.022e-15 |

D ENCODE transcription analysis

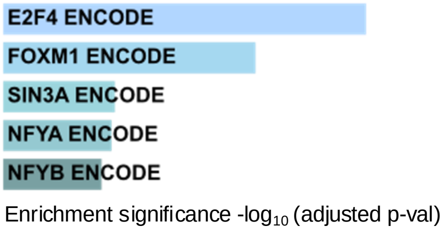


| ENCODE term | p-value | adj. p-value |
|---|---|---|
| E2F4 ENCODE | 4.635e-64 | 4.589e-62 |
| FOXM1 ENCODE | 9.170e-44 | 4.539e-42 |
| SIN3A ENCODE | 6.118e-18 | 2.019e-16 |
| NFYA ENCODE | 2.446e-17 | 6.054e-16 |
| NFYB ENCODE | 1.687e-15 | 3.339e-14 |

**Supplementary Figure S12: (A,B)** Bar plots showing the activity of significant ($p < 0.05$) TFs relative to PC12 cells treated with PI3Ki (buparlisib) *vs.* control (A) and combination (Combo) *vs.* control (B). The activity score was calculated using the WMEAN (weighted mean) method. Red (score > 0) indicates high activity, blue (score < 0) indicates low activity. **(C,D)** Bar plots show the top five results from the enrichment analysis performed using Enrichr with the ChEA2022 and ENCODE categories. On the right of the bar plots are displayed the p-values and adjusted p-values for each enriched term. Plots were generated using R software (v 4.4.0).

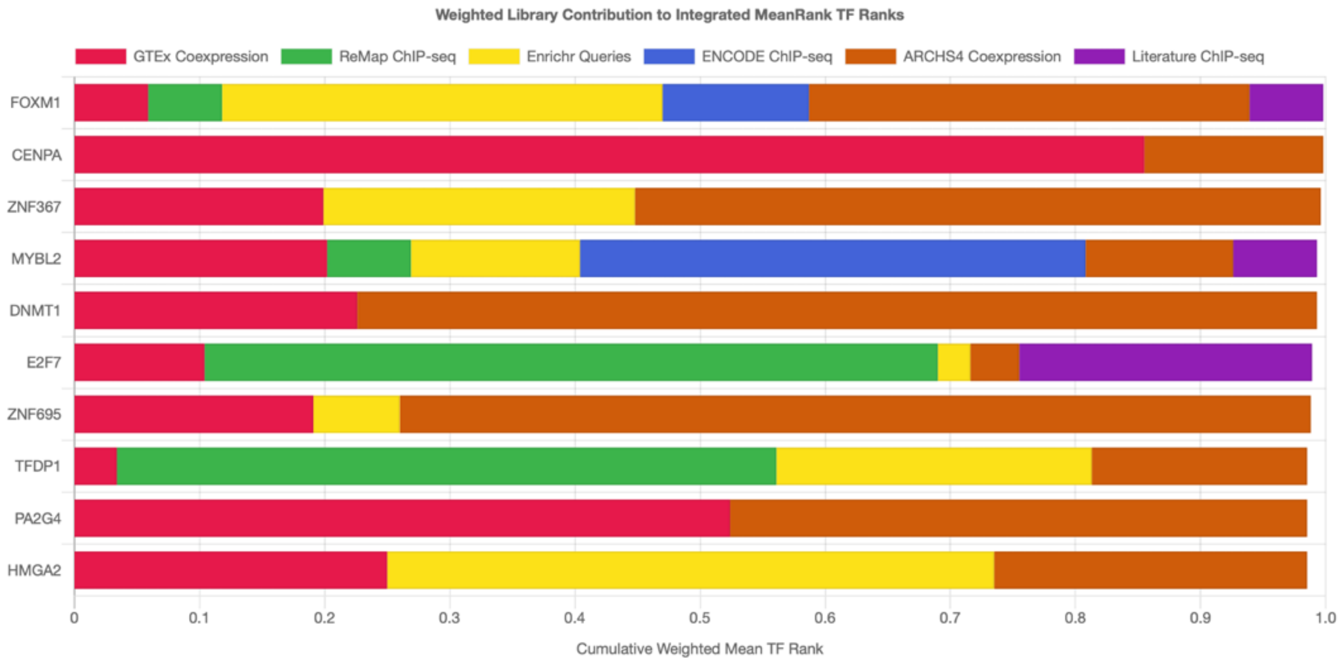


**Supplementary Figure S13.** Stacked bar plot indicating the enrichment of TFs regulating the genes uniquely downregulated in PC12 cells upon combination treatment, used as input in the ChEA3 tool. The different colors of the bars represent various curated databases containing ChIP-seq information. The x-axis indicates the cumulative weighted mean calculated considering each database.

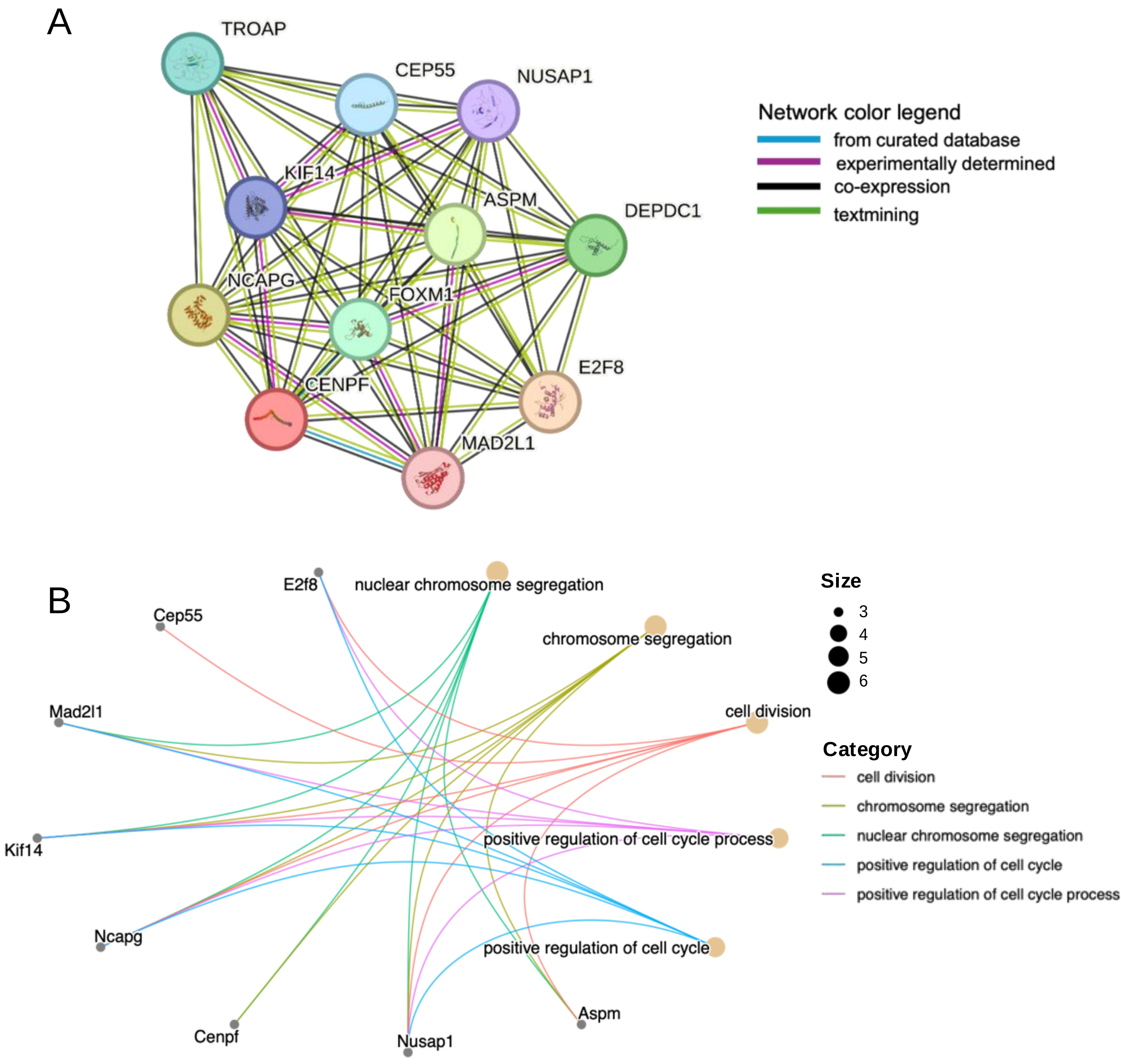


**Supplementary Figure S14**: **(A)** Interaction network graph, generated using STRING database, displaying the predicted and experimentally validated interactions among the selected proteins. Each node represents a protein, while lines indicate the associations among proteins, including physical interactions and functional relationships. The thickness and color of the edges reflect the confidence scores of these interactions, with stronger lines indicating higher confidence. **(B)** Circular net plot visualizing the relationships between genes and enriched pathways belonging to GOBP terms identified through enrichment analysis. In this network, genes are represented as circular nodes, and pathways or GO terms as larger nodes. Edges connect genes to the pathways they are involved in, allowing for the identification of key genes shared across multiple pathways. The size and color of the nodes reflect the significance of enrichment.

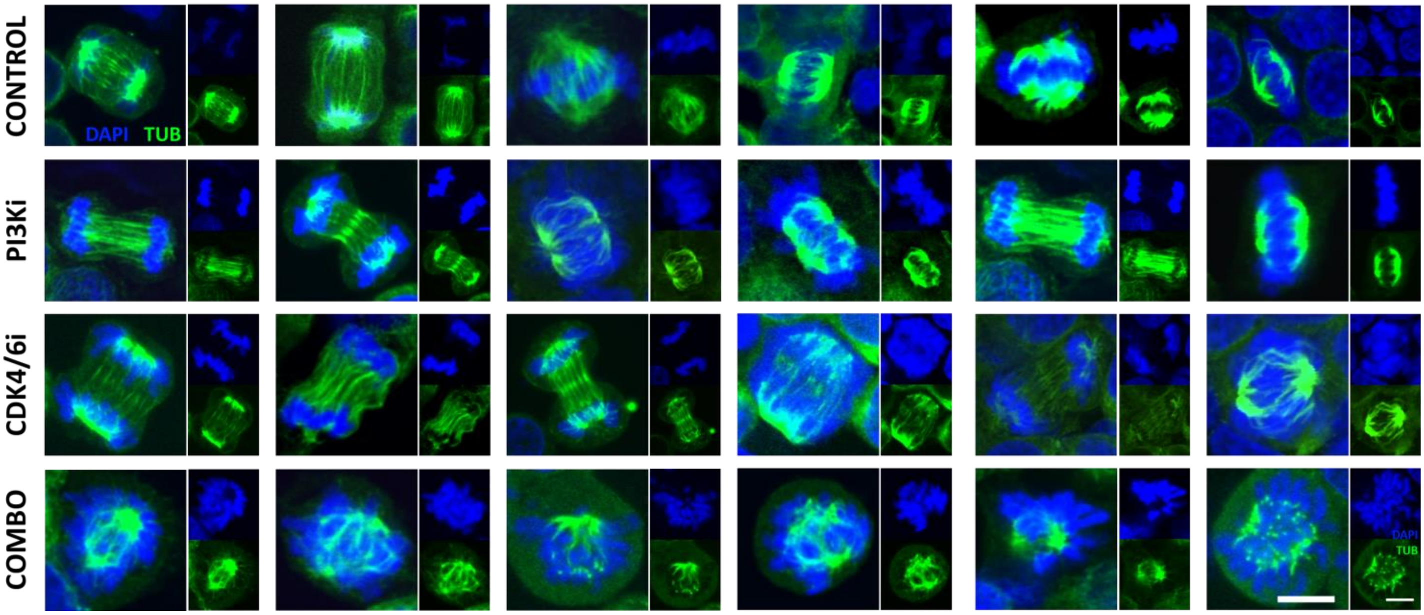


**Supplementary Figure S15.** Characterization of mitotic spindle in PC12 cells. Following individual and combined treatment with PI3Ki and CDK4/6i. PC12 cells cultured on Poly-L-lysine coated coverslips were either untreated or treated with one of 1µM PI3Ki or 4µM CDK4/6i or the combination for 24 h. Cells were then fixed, permeabilized and immunostained with antibody against α-tubulin (Green). Nuclei were counterstained with 4',6-diamidino-2-phenylindole (Blue). Mitotic cells were visualized with Leica TCS SP5 (× 63 magnification). For each condition, images of 30 mitotic cells were captured. 6 exemplary confocal images are shown. Scale bar (shown in the last montage) = 10µm.

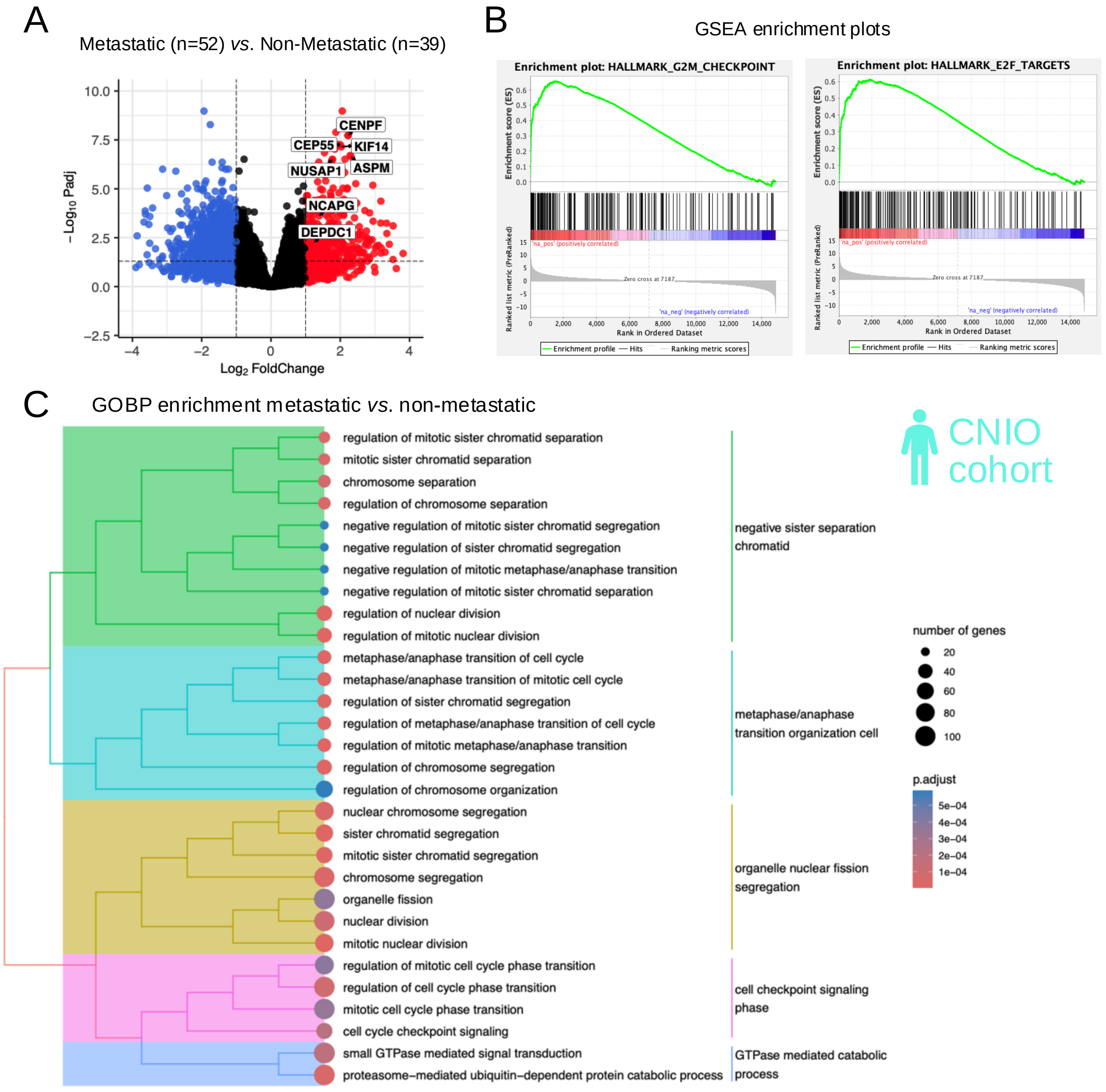


**Supplementary Figure S16.** **(A)** Volcano plot showcasing the distribution of DEGs between metastatic and non-metastatic samples in the CNIO cohort. Red points represent upregulated genes, blue points indicate downregulated genes, and black points mark non-significant changes. Significantly upregulated downstream targets of FOXM1 are highlighted. **(B)** GSEA plots illustrating the enrichment of upregulated genes in the G2M checkpoint and E2F targets hallmark pathways within the CNIO cohort. The X-axis represents genes ranked by expression levels, while the Y-axis shows the cumulative Enrichment Score (ES). The main curve depicts the ES trend, with a positive peak indicating enrichment of the gene set at the top of the ranked list. Vertical black ticks on the X-axis denote the positions of individual pathway genes within the ranked dataset, visualizing their density and distribution.**(C)** Tree plot visualizing enriched GOBP categories comparing metastatic versus non-metastatic samples, emphasizing significant biological processes grouped by functional similarities. Each node represents an enriched GO term, with node size corresponding to the number of associated genes. Proximity of nodes reflects related processes, and larger clusters indicate broader functional categories. The color gradient (red to blue tones) represents adjusted p-values, with red indicating higher significance. All plots were generated using R software (v4.4.0).

A

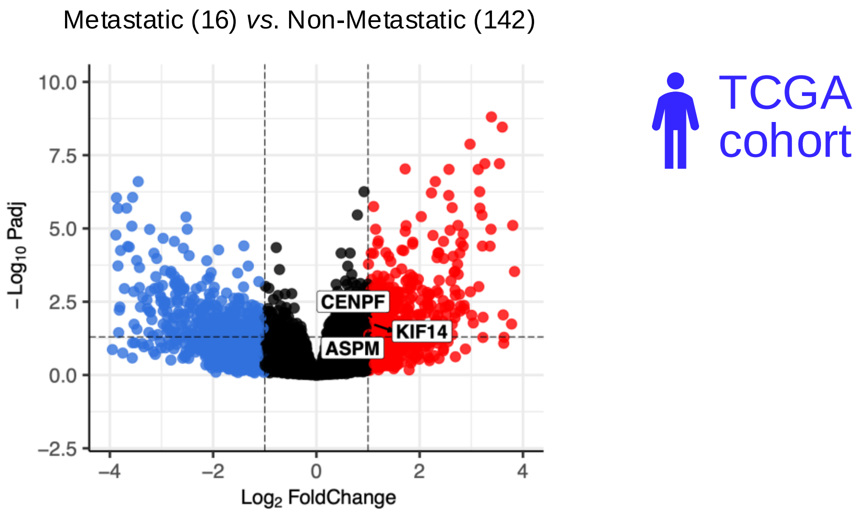


B

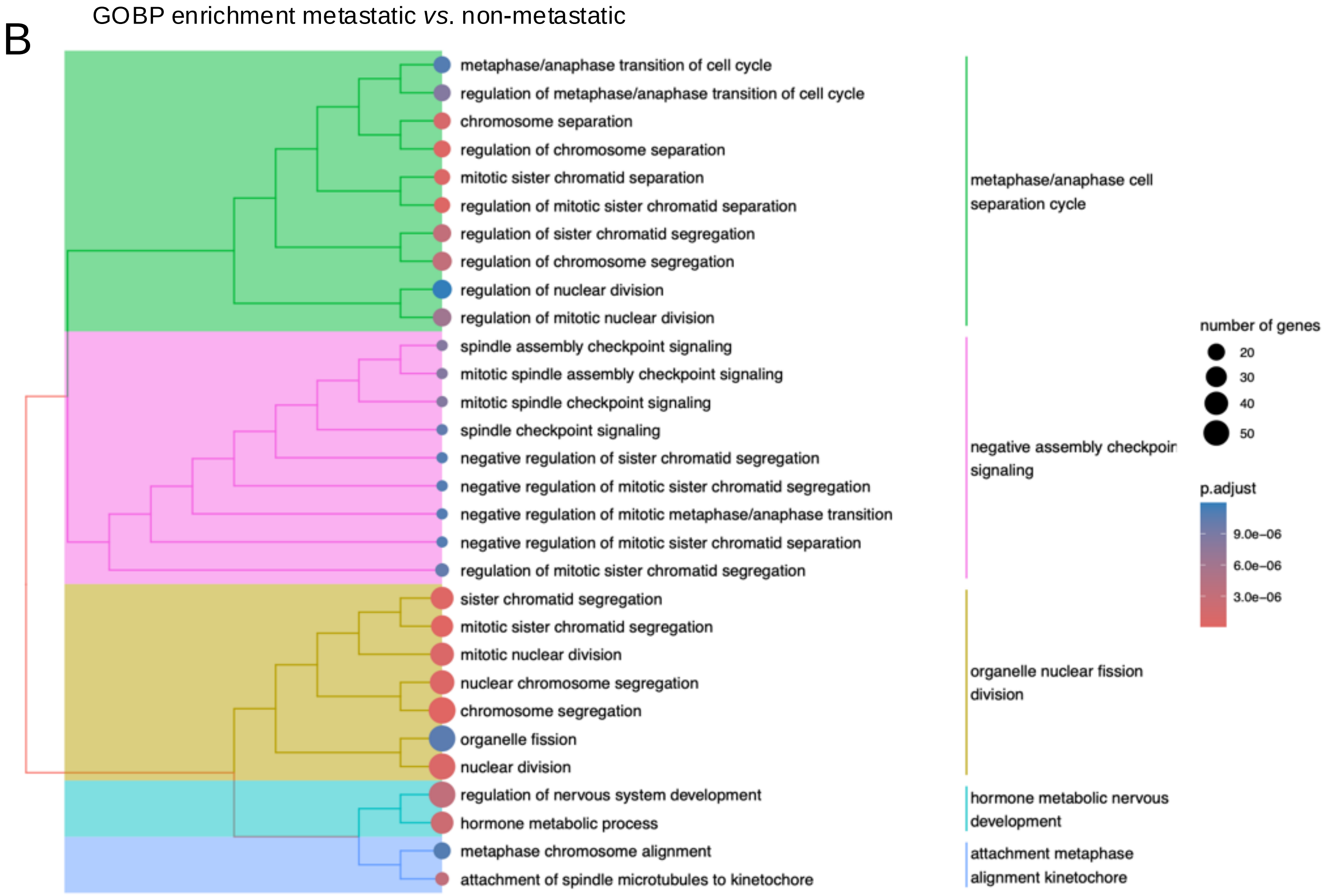


**Supplementary Figure S17. (A)** Volcano plot showcasing the distribution of DEGs between metastatic and non-metastatic samples in the CNIO cohort. Red points represent upregulated genes, blue points indicate downregulated genes, and black points mark non-significant changes. Significantly upregulated downstream targets of FOXM1 are highlighted. **(B)** Tree plot visualizing enriched GOBP categories comparing metastatic versus non-metastatic samples, emphasizing significant biological processes grouped by functional similarities. Each node represents an enriched GO term, with node size corresponding to the number of associated genes. Proximity of nodes reflects related processes, and larger clusters indicate broader functional categories. The color gradient (red to blue tones) represents adjusted p-values, with red indicating higher significance.

All plots were generated using R software (v4.4.0).

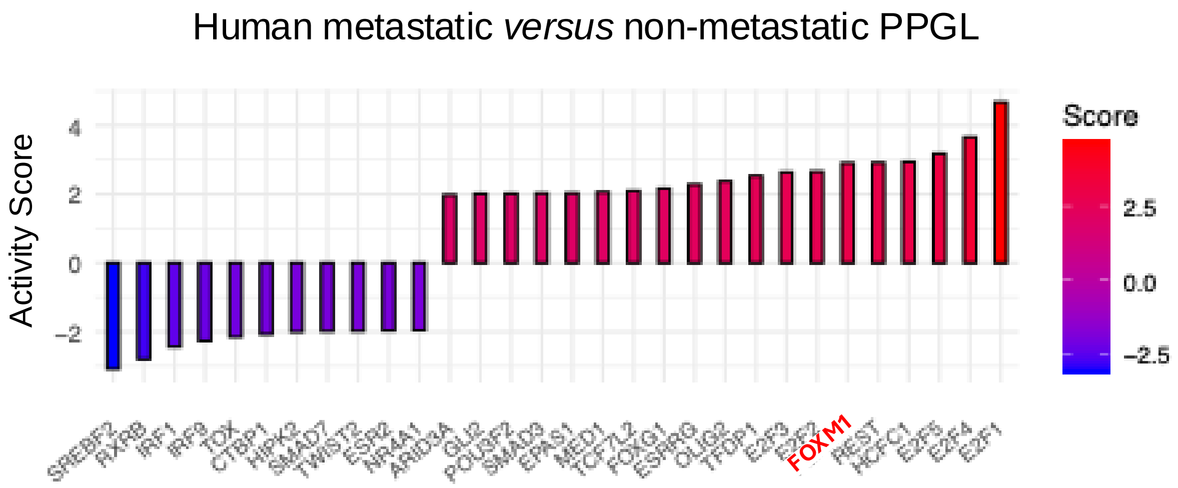


**Supplementary Figure S18.** Bar plots showing the activity of each transcription factor in the comparison between metastatic and non-metastatic samples in the CNIO cohort; the activity score was calculated using the WMEAN (weighted mean) method. Red (score > 0) indicates high activity, blue (score < 0) indicates low activity. The plot was generated using R software (v. 4.4.0).
